# Identification of antifungal agents AR-12 and Fosmanogepix as anti-*Trypanosoma cruzi* drugs through an enhanced fluorogenic β-galactosidase phenotypic screening assay

**DOI:** 10.1101/2025.10.01.679864

**Authors:** Mercedes Didier Garnham, Franco Agustín Agüero, Juan Carlos Ramírez, Fernán Agüero, Emir Salas-Sarduy

## Abstract

Phenotypic screening remains essential for identifying and characterizing bioactive compounds or their combinations against human parasitic pathogens. In the case of *Trypanosoma cruzi*, the etiological agent of Chagas disease, transgenic parasites expressing the reporter enzyme β-galactosidase have been extensively used to this end. Here, we replaced the traditional chromogenic substrate chlorophenol red-β-D-galactopyranoside (CPRG) with the fluorogenic 4-methylumbelliferyl-β-D-glucopyranoside (MUG) to derive a highly sensitive, continuous enzymatic assay to obtain a quantitative surrogate of parasite growth in *T. cruzi* cultures. The assay detects as few as 3×10^3^ trypomastigotes/well, tracks linearly with the parasite load in a two-order range (3×10^3^-2×10^5^ trypomastigotes/well), takes 1 hour, and has a similar cost per assay as its colorimetric counterpart. To demonstrate its convenience and versatility, we used this assay to estimate the half-maximal inhibitory concentration (IC_50_) of six emerging antifungal compounds, not targeting CYP-51 and novel for *T. cruzi*. Finally, the assay was adapted to a semi-automatic methodology and used to explore dual combinations of the active antifungals in the primary screening and with benznidazole. The multitarget compound AR-12 (IC_50_ = 1.9 μM) and the Gwt1 inhibitor Fosmanogepix (IC_50_ = 7.2 μM) resulted in *bona fide* hits, inhibiting parasite replication with only low-to-moderate toxicity on Vero host cells, thus suggesting potential for repurposing to Chagas disease.

## INTRODUCTION

Chagas disease is a debilitating, potentially fatal infection caused by the protozoan parasite *Trypanosoma cruzi*. This parasitosis, considered by the World Health Organization as a neglected tropical disease, remains endemic in 21 Latin American countries and is emerging as a global health threat, with 7-8 million people infected worldwide and no vaccine in the short term [1,2]. As the only two drugs in clinical use – benznidazole and nifurtimox – show limited efficacy in the chronic phase and a suboptimal safety profile [3–5], the identification of new therapeutic candidates (and their targets within the parasite) is critically important to populate the pipeline with promising drug candidates.

Different strategies, including drug repurposing [6,7], target-based [8–10], and phenotypic screenings of compound libraries [11–15], are commonly used to generate novel hits for clinical development against *T. cruzi*. In their early investigational stages, all these strategies converge in evaluating the antiparasitic activity of candidate compounds in *T. cruzi* cultures. One of the more frequently used assays for assessing anti-*T.cruzi* activity is based on transgenic parasites from the Tulahuen strain (Tul β-gal) constitutively expressing the β-galactosidase enzyme from *Escherichia coli* [16]. As mammalian cell hosts usually display negligible levels of endogenous β-galactosidase activity, this reporter enzyme is used as a proxy to estimate parasite-derived β-galactosidase concentration and, indirectly, the parasite load present in the culture [17].

As initially reported, this assay is based on the use of a galactopyranoside chromogenic substrate [18]. However, careful experimental conditions are needed to guarantee a direct linear translation from parasite load to β-galactosidase activity. From an enzymological perspective, it is necessary to monitor enzymatic activity under initial velocity conditions, ideally using a robust and sensitive continuous assay, allowing the unambiguous slope derivation from experimental progression curves. Further, the assay should also be able to discriminate between subtle differences in β-galactosidase levels (i.e., caused by different concentrations of test drugs in dose-response curves) and show a wide linear workable range (i.e., V_0_ vs. E_0_). Finally, high sensitivity at both high and low enzyme concentration ranges is highly desirable, as it would allow quantifying the early appearance of parasites (i.e., for wash-out experiments) and the evolution of parasite load over time (i.e., rate-of-kill assays) [17,19].

In our hands, we noted that the chromogenic substrate chlorophenol red-β-D-galactopyranoside (CPRG), traditionally used in this assay, is not well suited for the task. It exhibited low sensitivity, limited solubility under assay conditions, and erratic progression curves. Also, it required a long incubation time for color development before the single endpoint measurement. Even after a 4-hour incubation, absorbance signals in a typical experiment are around 0.4 and 0.15 for positive (infected) and negative (non-infected) controls, respectively, resulting in a signal-to-noise ratio of less than 3 and a dynamic range of 0.25 absorbance units.

To facilitate the phenotypic evaluation of new compound collections from several repurposing strategies, here we transformed the traditional endpoint chromogenic methodology into a more convenient, continuous, and semi-automatic fluorogenic assay by using 4-methylumbelliferyl-β-D-galactopyranoside (MUG) as substrate. Finally, we used this continuous methodology to assess the antiparasitic activity of six emerging antifungal drugs (Figure 1), not previously tested against *T. cruzi* and not targeting CYP-51. As a result, we propose the multitarget compound AR-12 and the Gwt1 inhibitor Fosmanogepix, exhibiting potencies in the low micromolar range and suitable selectivity indexes, as *bona fide* repurposing candidates for Chagas Disease.

**Figure 1.**
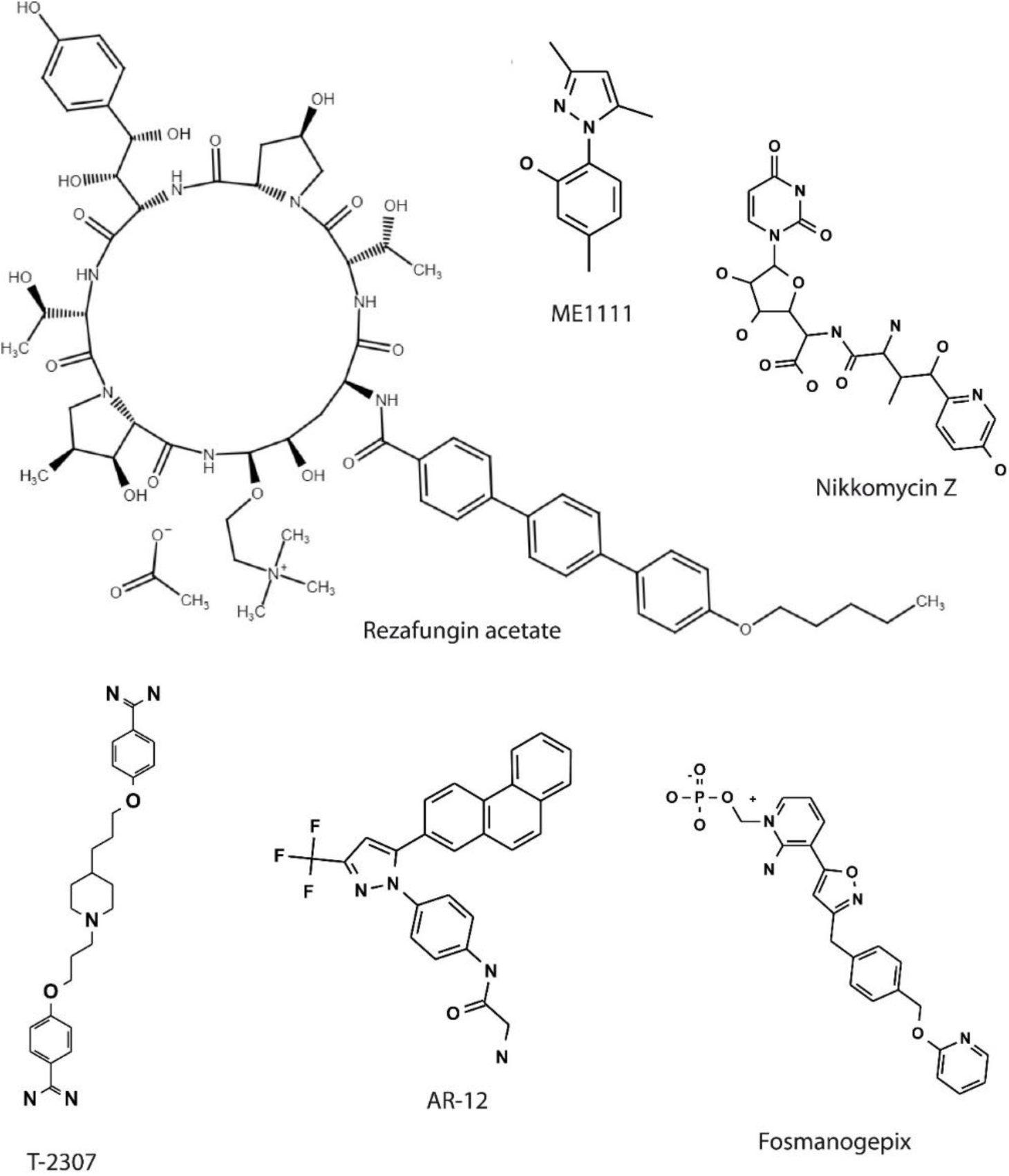
Chemical structures of the investigated antifungal compounds.

## RESULTS

### 1. Development of a continuous enzymatic β-galactosidase assay using MUG as fluorogenic substrate

As part of a diverse set of repurposing strategies in our laboratory, we recently evaluated small collections of synthetic compounds against *T. cruzi* in phenotypic screenings [20] by using Tul β-gal parasites and CPRG as chromogenic substrate [16]. Despite its simplicity and low cost, we found the long incubation, poor dynamic range, and endpoint nature of the assay troublesome for productivity and analytic purposes. We reasoned that these inconveniences derived from the low sensitivity of CPRG as a β-galactosidase substrate and hypothesized that its substitution by a fluorogenic alternative [21,22] would render a continuous assay better suited not only for primary screening and dose-response measurements, but also for more demanding applications such as rate-of-kill and wash-out experiments.

To test this idea, we evaluated the ability of MUG substrate to be hydrolyzed by parasite lysates containing β-galactosidase. Because our main interest was to design a continuous assay that would work properly at the sample composition resulting from cell rupture, we used PBS supplemented with 2% RPMI, 0.25% DMSO, and 0.5% NP-40, pH 7.4, as the initial activity buffer. The product of MUG hydrolysis, 4-methylumbelliferone, displays enhanced fluorescence in the basic pH range [23,24]. Thus, several protocols using MUG as a β-galactosidase substrate include the addition of a basic buffered solution (i.e., pH ≥ 10) as a final step before fluorescence measurement [25–30]. Because this would automatically transform the assay into an endpoint test, we eliminated this step in favor of continuous fluorescence monitoring at neutral pH in solid-black 384-well plates.

In contrast to control wells containing only activity buffer, the addition of MUG to wells containing parasite lysate as a β-galactosidase source produced a robust and linear increase of fluorescence over time, indicating the feasibility of continuous monitoring of the enzymatic hydrolysis of MUG at neutral pH (Figure 2A). A suitable working range for β-galactosidase was determined through the activity of two-fold serial dilutions of parasite lysate at a fixed substrate concentration of 10 μM. For a wide range of enzyme dilutions, progression curves remained linear for at least 40-60 minutes, a suitable time for recording initial velocity (V_0_), and the V_0_ vs. [E]_0_ plot displayed the expected linear behavior (Figure 2B). We next used a fixed β-galactosidase dilution (1/2,000) to analyze the dependence of V_0_ on MUG concentration in the range 0-10 μM (Figure 2C). Under these conditions, a linear behavior was observed, indicating low saturation rates for the enzyme even at the highest concentration tested (K_M_>>10 μM). Thus, we fixed the final MUG concentration in the assay to 10 μM to maximize enzyme activity while still avoiding the inner-filtering effect [31]. Finally, we explored the effect of DMSO (range: 0.5-2%) and NP-40 (range: 0.5-4%) on the continuous β-galactosidase assay, detecting activity drops of ∼ 20% at concentrations >2% for both components (Supplementary Figure S1). Under optimized conditions, the assay showed good overall performance, with a dynamic range (average dF/dt^C+^ - average dF/dt^C-^) > 5000 RFU/sec, a signal-to-noise ratio (average dF/dt^C+^/average dF/dt^C-^) > 9,000, coefficient of variation (CV) < 2%, and Z factor values in the range 0.7– 0.9.

**Figure 2.**
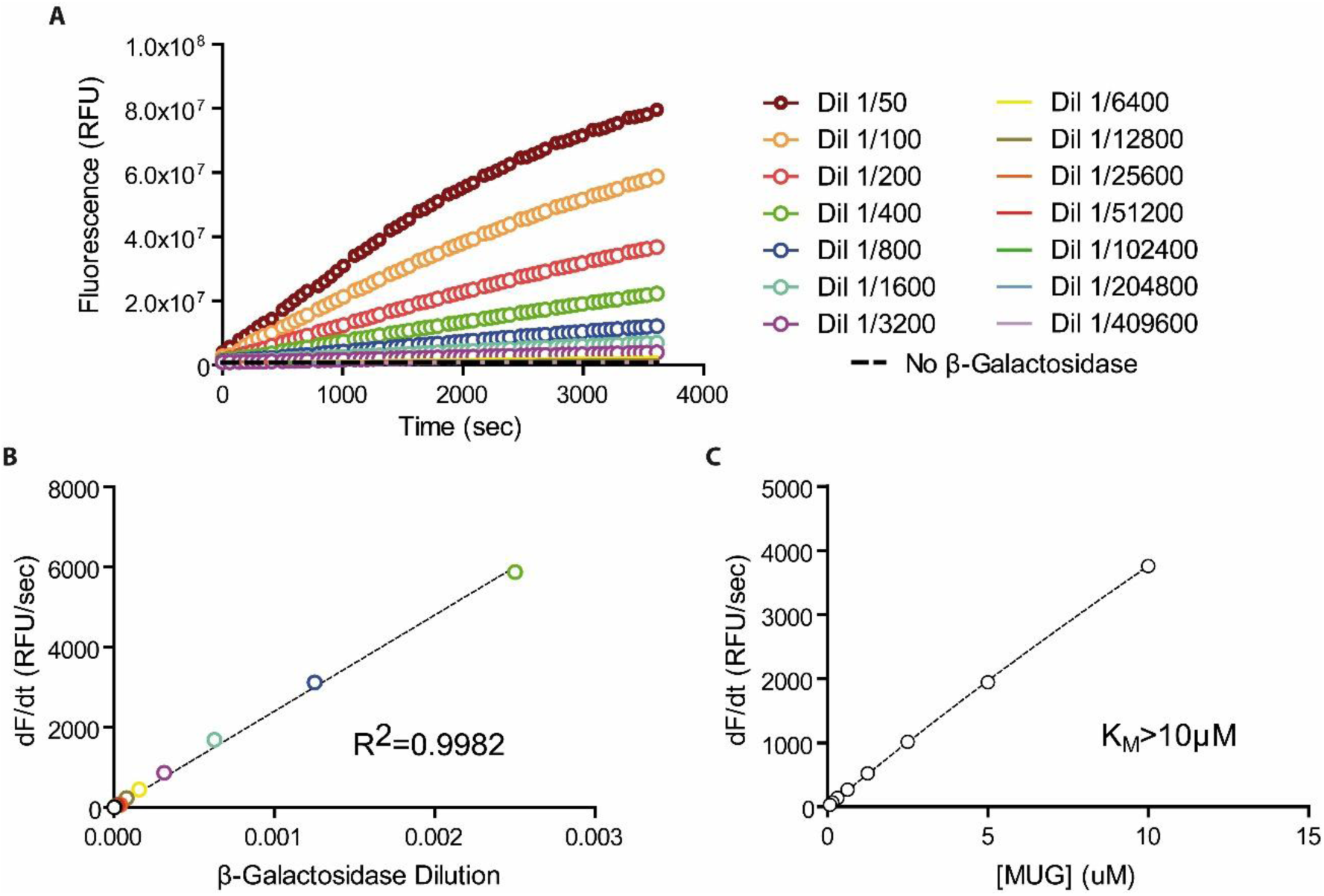
Characterization of the continuous β-galactosidase assay using MUG as substrate. A) Progression curves for the hydrolysis of MUG ([MUG]= 10 μM) by two-fold β-galactosidase dilutions (range 1/25 – 1/409,600). B) Plot of V_0_ vs. [E]_0_ for increasing β-galactosidase dilutions (range: 1/400 to 1/204,800) at a fixed substrate concentration ([MUG]_0_= 10 μM). Color code corresponds to the legend in panel A. D) Michaelis-Menten plot for β-galactosidase (enzyme dilution: 1/2,000).

### 2. Adaptation of the continuous fluorogenic methodology to fresh trypomastigote lysates

We next decided to evaluate the performance of the assay when applied to fresh cultures of Tul β-gal trypomastigotes. Initially, parasites were serially diluted and spiked into 96- well plates containing seeded Vero cells. After cell lysis, lysates were assayed in parallel for β-galactosidase activity using both substrates.

After 4 hours of incubation with CPRG, representative absorbances ranged from 0.11 to 0.33 (for 0 and 2×10^5^ trypomastigotes/well, respectively; Figure 3A). Absorbance tracked linearly the parasite number (R^2^=0.981) in the tested range, and analytical sensitivity was estimated to be 0.01 AU/1×10^4^ trypomastigotes. Despite the narrow dynamic range (0.21 AU) and low signal-to-noise ratio (2.8), the colorimetric assay showed Z factor values of 0.8-0.9 due to very low replicate variability.

**Figure 3.**
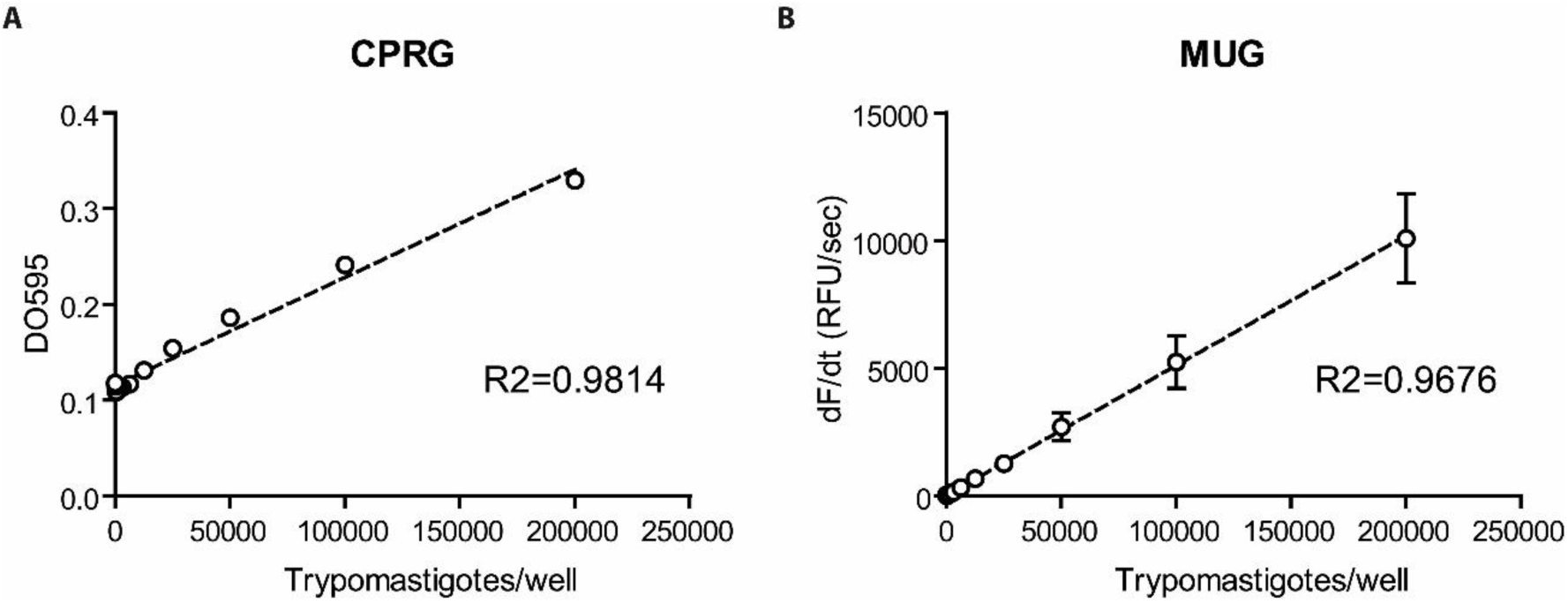
Linear regression analysis between signal and parasite load for both β-galactosidase assay variants. A) Plot of DO_595_ vs. trypomastigotes/well for the endpoint colorimetric β-galactosidase assay after a 4-hour incubation with CPRG substrate ([CPRG]= 50 μM). B) Plot of dF/dt vs. trypomastigotes/well for the continuous fluorometric β-galactosidase assay. Linear regression R^2^ values are indicated. Each point represents the average of four replicates.

Interestingly, progression curves generated with MUG were not linear in their initial section and steeped gradually until reaching a constant steady-state slope (Supplementary Figure S2). This behavior, only noticeable by the continuous nature of this assay, might arise from a combination of causes, including temperature equilibration, slow completion of parasite lysis, and substrate-induced renaturation of partially unfolded/catalytically defective enzyme populations. Steady-state slopes were obtained after approximately 20-25 minutes and remained constant for at least 60 more minutes (Supplementary Figure S3). These slopes correlated linearly with trypomastigote numbers in the complete range (R^2^=0.968) (Figure 3B), and the assay displayed an analytical sensitivity of 507 RFU*sec^-^/1×10^4^ trypomastigotes. In contrast with the endpoint assay, typical quantifications with MUG exhibited dynamic ranges > 10,000 RFU*sec^-^ and signal-to-noise ratios > 200. Typical Z factor values for the continuous fluorogenic methodology were 0.8-0.9, like those obtained with CPRG. Of note, the lowest number of parasites that caused signals above that of the negative control + 3SD (Vero cells with no spiked trypomastigotes) were 12,500 and 3,125 for CPRG and MUG, respectively, indicating that the latter assay displays an improved detection limit.

### 3. Adaptation of the continuous methodology to intracellular amastigotes by using the reference drug benznidazole

We then extended the continuous fluorogenic methodology to quantify intracellular Tul β-gal amastigotes. Considering the significant technical challenge of using a spike-in strategy in this case, we thought to apply increasing concentrations of the reference drug benznidazole (BNZ) to infected cells to gradually obtain a decreasing number of intracellular amastigotes and quantify β-galactosidase activity by both methodologies in the same experiment.

Table I summarizes the performance parameters for both assays in three independent experiments. As previously noted, small dynamic ranges, signal-to-noise ratios, and replicate variabilities were observed after a 4-hour incubation with CPRG. In contrast, the continuous quantification of β-galactosidase activity with MUG led to dynamic ranges > 8,000 RFU*sec^-^ and signal-to-noise ratios > 60. As observed in the previous experiments, replicates in the fluorogenic assay showed moderate variability. Despite these differences, both methodologies consistently showed highly similar Z factor values (≥ 0.79), indicating good overall performance. More importantly, both methodologies sensed decreasing levels of β-galactosidase activity as the BNZ concentration increased in the assay and derived highly similar dose-response curves for BNZ (Figure 4 and Supplementary Table SI). The average IC_50_ values for the endpoint and the continuous assays were estimated to be 0.796 ± 0.132 μM and 0.754 ± 0.193 μM, respectively, showing no significant statistical differences.

**Figure 4.**
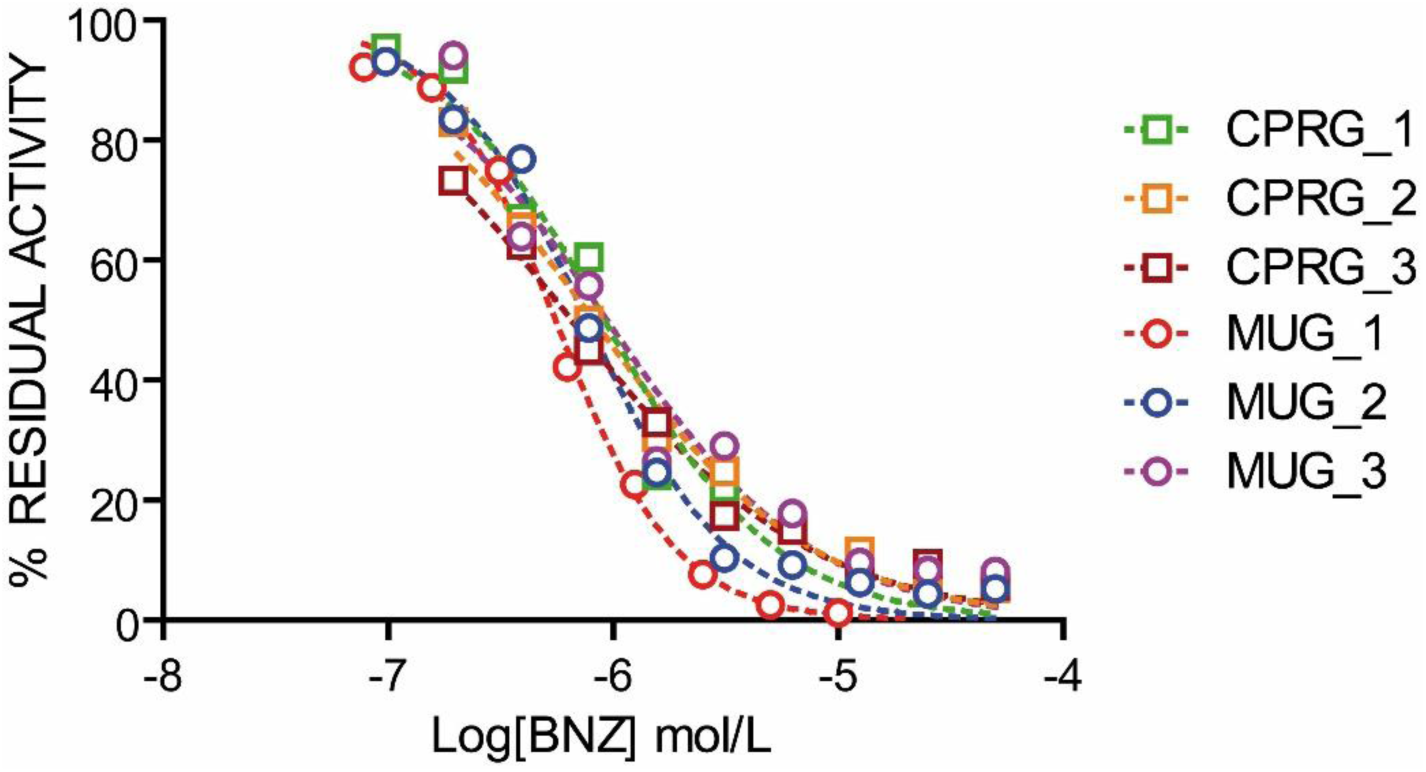
Representative dose-response curves for the reference drug benznidazole obtained using both methodologies. The endpoint chromogenic assay using CPRG as substrate is represented by open squares and the continuous fluorogenic methodology using MUG substrate by open circles. Three independent experiments are represented.

**Table I.** Comparison of assay performance in the detection of β-galactosidase activity in intracellular amastigotes.

| Assay | Exp | C-<br>Average ± SD | C+<br>Average ± SD | Dynamic<br>range | Signal-to-<br>noise ratio | Z factor |
| --- | --- | --- | --- | --- | --- | --- |
| Chromogenic<br>(Endpoint) | 1 | 0.1237 ±<br>0.0032 AU | 0.467 ± 0.0084<br>AU | 0.3432 AU | 3.773 | 0.898 |
| | 2 | 0.1151 ±<br>0.0030 AU | 0.3197 ± 0.0109<br>AU | 0.2046 AU | 2.778 | 0.797 |
| | 3 | 0.1069 ±<br>0.0011 AU | 0.3006 ± 0.0077<br>AU | 0.1937 AU | 2.813 | 0.864 |
| Fluorogenic<br>(Continuous) | 1 | 210.83 ± 31.94<br>RFU/sec | 13,598.89 ±<br>526.69 RFU/sec | 13,388<br>RFU/sec | 64.503 | 0.875 |
| | 2 | 100.19 ± 5.00<br>RFU/sec | 8,373.63 ±<br>550.68 RFU/sec | 8,273<br>RFU/sec | 83.576 | 0.799 |
| | 3 | 123.18 ± 8.901<br>RFU/sec | 10,178.37 ±<br>260.97 RFU/sec | 10,055<br>RFU/sec | 82.631 | 0.919 |

To evaluate whether the increased inter-replicate variability observed with MUG was due to actual biological differences or introduced methodological artifacts, we repeated the previous experiment using a highly sensitive quantitative real-time PCR assay [32] to estimate *T. cruzi* DNA loads in the lysates, in parallel with CPRG and MUG enzymatic assays. The qPCR standard curve (range: 10^5^ to 10 parasite equivalents/mL, R^2^=0.999, Efficiency =0.95) and a summary of Cts and the estimated parasite equivalent/mL in each sample replicate are shown in Supplementary Figure S4A and Supplementary Table II, respectively. Independent of the enzymatic methodology used, β-galactosidase activity strongly correlated with *T. cruzi* DNA loads (Figure 5A, B, and Supplementary Figure S4C, D), with estimated Pearson correlation coefficient (r) values of 0.9917 and 0.9723 for colorimetric (CPRG) and fluorometric (MUG) assays, respectively. Considering replicate variability, the coefficients of variation from the fluorogenic assay showed fair correlation with those from qPCR (Figure 5D and Supplementary Figure S4F), with Pearson r values of 0.8322 and 0.8199 for parasite equivalents/mL and Ct, respectively. On the contrary, CV values from the CPRG assay showed no correlation to those from the qPCR assay (Pearson r values of 0.02303 and -0.02732) (Figure 5C and Supplementary Figure S4E). These results indicate that the chromogenic assay underestimates the existing biological differences between replicates, probably due to the low sensitivity of the CPRG substrate.

**Figure 5.**
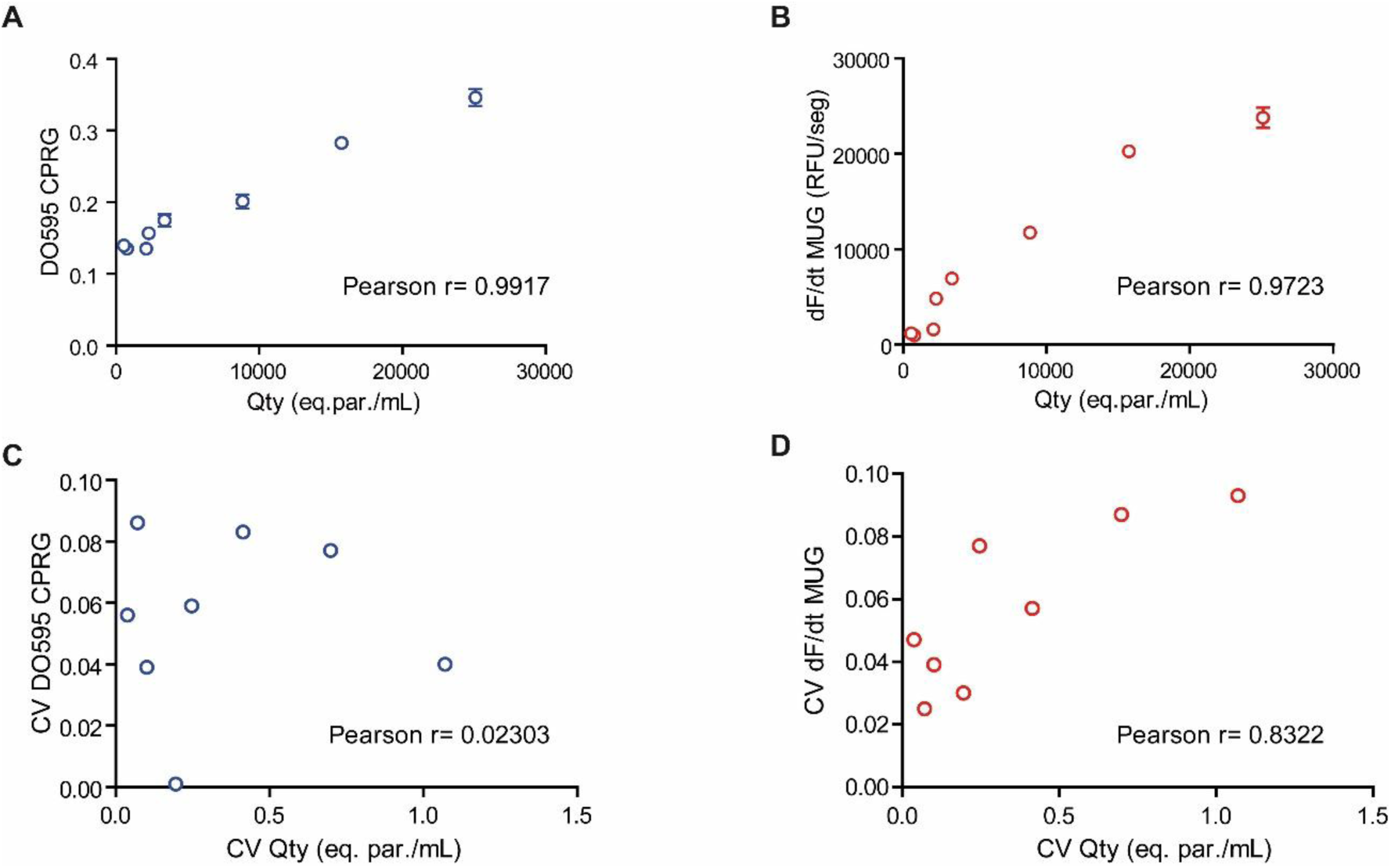
Correlation analysis between β-galactosidase activity and the amount of T. cruzi DNA estimated by qPCR. A) Endpoint assay (DO595) vs. qPCR (parasite equivalents/mL). B) Continuous assay (dF/dt) vs. qPCR (parasite equivalents/mL). Correlation analysis between the coefficient of variations (CV) from enzymatic methodologies and qPCR. C) CV of endpoint assay (CV DO595) vs. CV of qPCR (CV parasite equivalents/mL). D) CV of Continuous assay (CV dF/dt) vs. CV of qPCR (CV parasite equivalents/mL). In all cases, the estimated Pearson r values are indicated.

Overall, these results indicate that the continuous methodology derived here for the estimation of β-galactosidase activity can be safely adopted for the dose-response evaluation of investigational compounds in *T. cruzi* phenotypic screenings, with the additional benefits of reduced time consumption, augmented sensitivity, and realistic detection of inter-replicate variability.

### 4. Phenotypic screening of six emerging antifungal drugs using the continuous methodology

In a previous work, we identified nearly 3,000 shared drug-target associations between the yeast *Saccharomyces cerevisiae* and *T. cruzi*, encompassing more than 2,000 candidate compounds with potential bioactivity against the parasite [20]. A novelty analysis from the literature indicated that, from the 210 most promising candidates, 31 had been tested active against *T. cruzi* or other trypanosomatids in previous studies, meaning a 14.8% success rate in the worst-case scenario. These findings, in addition to the well-demonstrated anti-trypanosomatid activity of numerous antifungal drugs [33–39], encouraged us to evaluate in phenotypic screenings emerging antifungals not previously tested against *T. cruzi*.

For this study, we prioritized i) antifungal drugs that entered clinical trials in recent years, that were also ii) commercially available at affordable prices, and iii) had no reports of previous testing on *T. cruzi*. We decided to include emerging antifungals independently of their target or mechanism of action, except for CYP-51 inhibitors, which were excluded due to very high levels of parasite relapse in clinical trials [40,41]. The final list included the following six drugs: AR-12 (OSU-03012), ME1111, Fosmanogepix (E1210), Rezafungin acetate, T-2307, and Nikkomycin Z (Figure 1).

The antifungal drugs (and BNZ as positive control) were tested in *T. cruzi* phenotypic screening using the continuous fluorogenic methodology validated in the previous section. Rezafungin acetate, T-2307, and Nikkomycin Z showed negligible antiparasitic activity in culture (Supplementary Figure S5). On the other hand, ME1111, Fosmanogepix, and AR-12 exhibited reproducible anti-*T. cruzi* activity, displaying typical dose-response curves and estimated IC_50_ values of 19.69 μM, 7.16 μM, and 1.91 μM, respectively (Figure 6 and Table II). Next, we evaluated the cytotoxicity of the six antifungal drugs on Vero cells, using resazurin as a cell-permeable redox indicator to monitor culture viability [42]. At the highest concentration tested, five compounds (ME1111, Fosmanogepix, Rezafungin acetate, T-2307, and Nikkomycin Z) displayed only negligible cytotoxicity on host cells. In agreement with previous reports [43], AR-12 showed modest but detectable cytotoxicity on Vero cells (CC_50_=20.75 μM) and a selectivity index of 10.86, acceptable for its initial repurposing.

**Figure 6.**
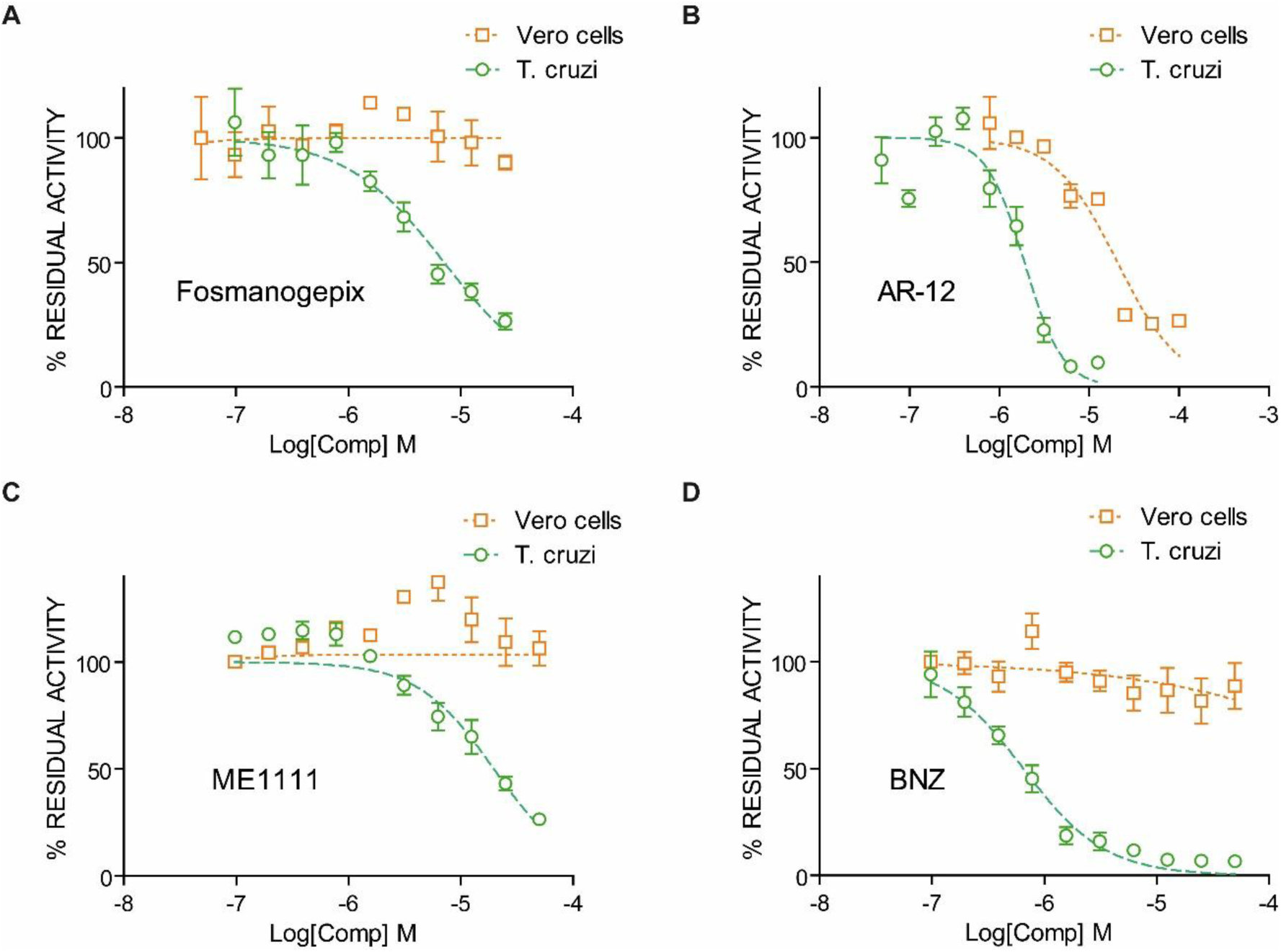
Dose-response curves for the antiparasitic and cytotoxic activities of active antifungal compounds. Antiparasitic activity against T. cruzi is represented with green circles, and the cytotoxic activity on Vero cells with orange squares. A) Fosmanogepix. B) AR-12. C) ME1111. D) Benznidazole (positive control). Each data point represents the average of six replicates.

**Table II.** Bioactivity summary of antifungal compounds.

| Drug | IC<sub>50</sub> (μM) on *T. cruzi*<br>(95% Confidence Interval) | CC<sub>50</sub> (μM) on Vero Cells<br>(95% Confidence Interval) | IC<sub>50</sub> (μM) on β-galactosidase | Selectivity Index |
| --- | --- | --- | --- | --- |
| AR-12 | **1.910**<br>(1.586 – 2.299) | **20.75**<br>(16.44 – 26.19) | > 100 | > 10.864 |
| Fosmanogepix | **7.155**<br>(5.383 – 9.511) | > 25 | > 25 | > 3.494 |
| ME1111 | **19.69**<br>(14.95 – 25.93) | > 50 | > 50 | > 2.539 |
| Rezafungin Acetate | > 50 | > 50 | > 50 | 1 |
| T-2307 | > 12.5 | > 12.5 | > 12.5 | 1 |
| Nikkomycin Z | > 50 | > 50 | > 50 | 1 |

Considering that inhibitors of the reporter enzyme would confound the interpretation of phenotypic screenings by mimicking truly active compounds able to block parasite replication, we tested the six antifungal drugs and isopropyl β-D-1-thiogalactopyranoside (IPTG, positive control) in a β-galactosidase enzymatic triage, using the fluorogenic assay developed in section 1. While, as expected, IPTG inhibited the reporter enzyme in a dose-dependent manner, the antifungal compounds showed no inhibitory activity on β-galactosidase in the range of concentrations tested (Supplementary Figure S6).

Overall, three of the six antifungal drugs tested were active against *T. cruzi*, displaying IC_50_ values below 25 μM and non-to-moderate cytotoxicity on Vero cells. Of note, none inhibited β-galactosidase in a triaging enzymatic assay, indicating that they promote actual dose-response reduction in parasite load in culture. Considering that AR-12 and Fosmanogepix display IC_50_ values in the low micromolar range and suitable selectivity indexes, they excel as promising chemical starting points for further optimization and SAR studies.

### 5. Partial automation of the methodology to facilitate the evaluation of drugs and their combinations

To facilitate and accelerate compound testing at medium-throughput, we adapted the continuous fluorogenic methodology into a semi-automatic pipeline that uses an affordable automated liquid handling robot (OpenTrons OT-2). Except for steps requiring a biosafety cabinet because of sterility or biological risk (cell seed, cell infection, and compound addition to cell cultures), all other individual operations of the assay (preparation of serial dilutions and compound combinations, addition of lysis buffer to the plates, cell lysate homogenization and transference to solid-black 384-well plates for fluorescence measurement, and addition of MUG substrate) were performed by the OT-2 robot via *ad-hoc* Python scripts, available in a GitHub code repository (https://github.com/trypanosomatics/OT2-protocols-for-antifungal-essays).

We validated the semi-automatic pipeline by assessing dual combinations of the active antifungals Fosmanogepix and AR-12, as well as their combinations with BNZ (Supplementary Figure S7). After the continuous quantification of β-galactosidase activity, resultant effect matrices were analyzed with SinergyFinder [44,45] and SynergyFinder Plus [46]. For the three combinations, synergy distribution maps showed complex landscapes alternating antagonism and synergy regions (Figure 7), independently of the scoring model used (Loewe, Bliss, HSA, and ZIP). Table III summarizes specific dose combinations displaying synergy for all scoring models and strong antiparasitic effect (inhibition of parasite replication ≥ 80%), as indicated by the Synergy Barometer tool [46]. In particular, the combinations of both antifungals with the reference drug BNZ showed promising results, suggesting potential for combined therapy regimens. Interestingly, at specific doses (see Table III for details), the combination of AR-12 and Fosmanogepix also displayed high therapeutic potential. Importantly, these antifungal dose combinations showed negligible cytotoxicity on Vero cells (cell viabilities of 94.95% and 96.7%, respectively), indicating an alternative therapeutic route for future evaluation in animal models.

**Figure 7.**
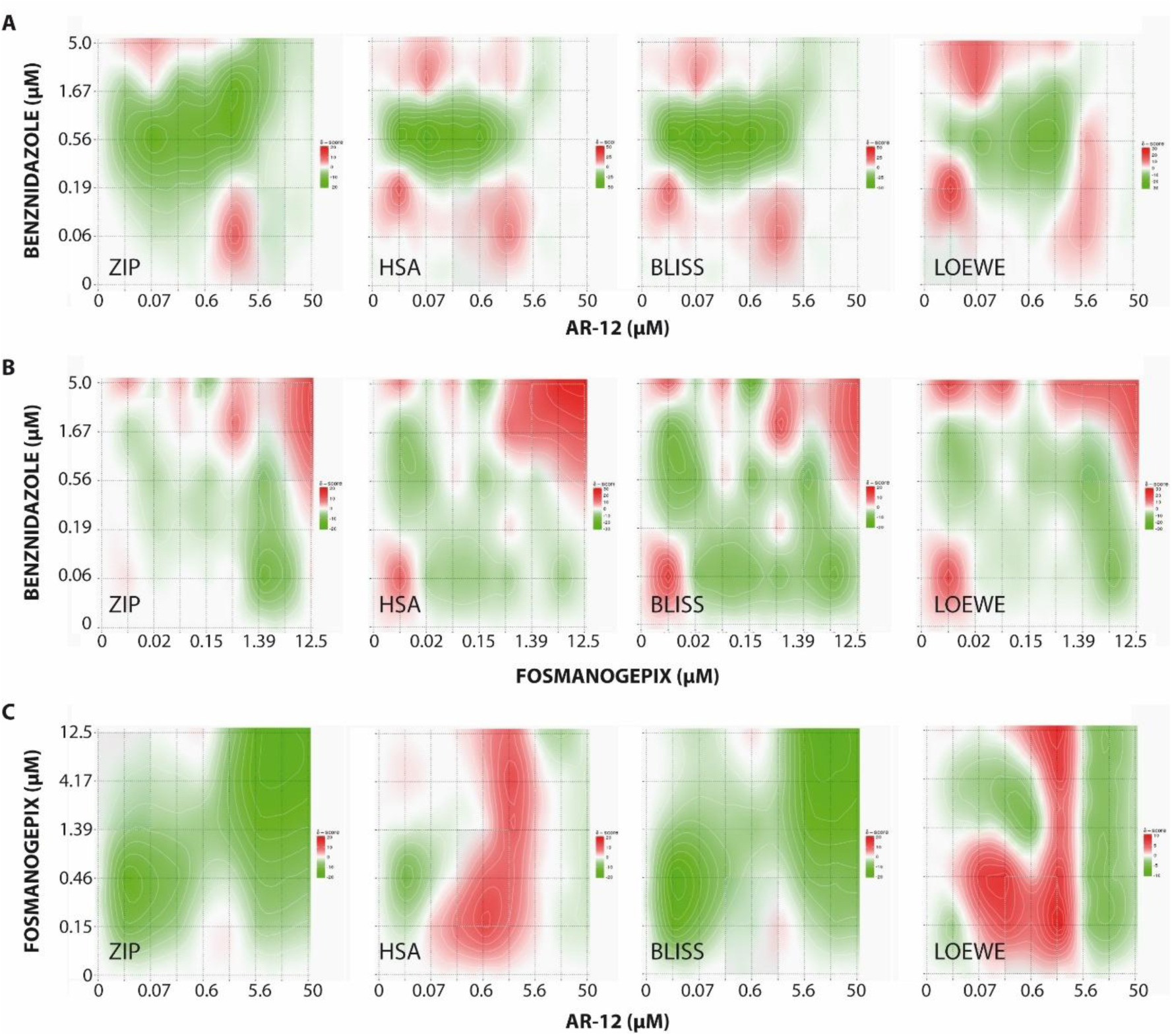
Synergy distribution maps for the combinations of the repurposed antifungal drugs with benznidazole. Four scoring models - ZIP, HSA, Bliss, and Loewe - were used to identify specific concentrations of drugs that, combined, resulted in antiparasitic effects significantly higher than those elicited by individual drugs at the same concentrations (synergy). A) AR-12/benznidazole. B) Fosmanogepix/benznidazole. C) AR-12/Fosmanogepix. The final concentration of each drug in the combination is indicated in the corresponding axis. For each combination, drug concentrations resulting in synergism are represented in red, while those resulting in antagonism are represented in green. Drug concentrations resulting in additive effects are represented in white.

**Table III.** Summary of selected dose combinations showing synergy for all the scoring models and a strong antiparasitic effect.

| Drug 1/Drug 2 | Dose drug 1 (μM) | Dose drug 2 (μM) | Calculated synergy scores for different models** |  |  |  | Observed Antiparasitic effect (%)* |
| --- | --- | --- | --- | --- | --- | --- | --- |
|  |  |  | ZIP | Loewe | Bliss | HSA |  |
| BNZ/AR-12 | 5.0 | 0.069 | 6.43 | 8.476 | 3.231 | 3.836 | 85.01 |
| BNZ/<br>Fosmanogepix | 5.0 | 12.5 | 9.939 | 14.2 | 9.303 | 24.46 | 97.01 |
|  | 1.667 | 12.5 | 13.78 | 10.02 | 11.75 | 17.71 | 90.26 |
|  | 5.0 | 4.167 | 4.483 | 18.33 | 5.432 | 26.97 | 87.96 |
| Fosmanogepix<br>/AR-12 | 12.5 | 0.6173 | 1.198 | 5.709 | 1.552 | 4.673 | 88.14 |
|  | 0.1543 | 1.852 | 0.4987 | 5.805 | 1.951 | 6.772 | 82.62 |

\* Only dose combinations displaying inhibition of parasite replication ≥ 80% are shown.
\*\* Values represent synergy scores, which indicate the degree of interaction predicted by each model for the drug combination at the indicated concentrations. The higher the score, the higher the synergy predicted by the model, meaning that the observed (antiparasitic) effect exceeds the expected additive response calculated by the model. A strong synergy should be concluded for a drug combination at specific concentrations if positive synergy scores are obtained from all the models [46]. Conversely, negative synergy scores indicate antagonism; observed effects are lower than the expected additive response. Synergy score values near zero indicate no interaction (additivity) between both drugs at the specific concentrations; i.e., no significant differences exist between the observed and calculated response effects.

## DISCUSSION

More than a century after its discovery, Chagas disease remains the most important tropical disease in the Americas and a major global health concern due to its continuous spreading into new regions. Although there have been remarkable advances in drug research for Chagas disease in recent years and several candidates have progressed to clinical trials, no new drug has yet reached therapeutic use, and the development pipeline remains poorly populated [47,48]. In this scenario, accurate and robust methodologies for phenotypic screening are critically important, as they can contribute to facilitate and accelerate the identification and characterization of promising candidates acting through novel mechanisms of action.

By substituting the chromogenic β-galactosidase substrate CPRG by the fluorogenic MUG, we transformed the traditional 4-hour long endpoint assay into a continuous enzymatic assay that can be completed in less than 60 minutes. In addition to time saving, the fluorogenic assay showed superior sensitivity, limit of detection, signal-to-noise ratio, and dynamic range than its chromogenic predecessor. Of note, the fluorogenic continuous assay is sensitive enough to capture the subtle differences that occur among biological replicates (unveiled by qPCR as reference method), which are underestimated by the endpoint assay. Given that fluorescence measurements require black plates to minimize well-to-well crosstalk, the continuous assay involves an additional transfer step from culture (transparent) to reading plates. This slightly increases the operational complexity of the assay and may contribute to higher replicate variability. These two factors may explain, at least partially, why the developed fluorometric assay displays Z factor values similar to those of the chromogenic one, despite its wider dynamic range. Importantly, the continuous nature of the fluorogenic assay offers a significant advantage over the previous endpoint assay, providing superior accuracy and reproducibility in slope estimations due to the multiplicity of experimental measurements taken during the initial velocity phase [49,50]. This is particularly true in cases like the CPRG-based chromogenic β-galactosidase assay, where extremely long experimental times are required, increasing chances of linearity loss for the enzymatic reaction by diverse factors, including: (i) β-galactosidase inactivation/degradation, (ii) substrate depletion, (iii) sample concentration by evaporation, and (iv) aggregates formation, among others.

The developed fluorogenic assay is highly versatile in several ways. First, it can be effortlessly transformed into an endpoint assay, if the number of tested compounds in primary screening (single dose) is very high (>> 384) and/or maximal accuracy is not required. As in the case of the continuous assay, an incubation time of at least 25 minutes is mandatory to ensure that measurement(s) match the steady-state phase. In this endpoint schema, the addition of a basic buffer (pH ≥ 10) could also be implemented before fluorescence reading, with the benefit of enhanced sensitivity [30]. For dose-response curves, continuous fluorescent monitoring is strongly recommended, although the recording time can be reduced from 60 to 20 minutes in the steady-state phase. While automation confers important functional advantages to the developed methodology, including reduced human error rates, increased consistency, and higher speed/throughput capabilities, this assay can be run manually if an automatic liquid handling system is not available. Finally, the robustness of the reporter β-galactosidase enzyme and MUG substrate would allow for the modification of assay conditions (buffer composition, detergent type/concentration, DMSO addition, culture media, etc.) to accommodate different host cells or investigational compounds with only modest impact on sensitivity.

During the first 20 minutes of the continuous assay, a consistent loss of linearity is observed in almost all wells. This loss is self-corrected over time. Although the exact nature of this phenomenon is yet to be determined, we opted to include an unmonitored (“blind”) incubation period and displace fluorescence monitoring to the steady-state phase to facilitate the analysis of kinetic data for slope estimation. This poses a limitation to the assay, increasing total time and reducing throughput. Another practical limitation of our experimental design is the need for an additional transfer step before fluorescence measurement. However, this can be conveniently solved by using solid-black culture plates (e.g., Greiner Bio-One 655086 or similar) instead of standard transparent ones, allowing not only direct plate reading, but also the addition of MUG substrate along with the lysis buffer. This would also permit to eliminate the substrate addition step and unify lysis and “blind” incubations into one, reducing further the duration, the complexity, and inter-replicate variability of the assay. Finally, we speculate that the sensitivity and limit of detection of the continuous assay might be further improved by changing the excitation wavelength to 320-330 nm, as this would increase the fluorescence signal of the 4-MU product by nearly two-fold at neutral pH [24,51]. Such an increase in sensitivity would be of great interest to derive more demanding applications from this enzymatic methodology, extending its use to characterizing the rate-of-kill and the cidality of investigational compounds in wash-out experiments. Improved fluorogenic methodologies are currently under development in our laboratory for both applications.

Recently, a Dm28c *T. cruzi* strain expressing *E. coli* β-galactosidase as reporter enzyme was developed [52]. The continuous fluorogenic methodology described here could also be used with this strain, thus increasing the confidence of candidate characterization against multiple strains from different DTUs or facilitating drug screening in other life cycle stages of *T. cruzi* [53]. This enzymatic strategy could also fuel the identification of new drug candidates for other diseases caused by trypanosomatids, as diverse transgenic strains of *Leishmania sp.* expressing β-galactosidase have been reported and are being used for this purpose [54,55]. In addition to *E. coli* β-galactosidase, different luciferase variants have been used as reporter genes for developing *T. cruzi* bioassays for compound screening [56–58]. A novel Dm28c transgenic strain expressing luciferase was recently created, allowing for a sensitive bioluminescent reporter assay for drug evaluation in 96-well plate format [59]. This system, based on a red-shifted luciferase variant from *Photinus pyralis* [60], displays a dynamic range of four orders of magnitude (10^3^-10^7^ parasites), an estimated limit of detection of ∼200 amastigotes/well, and a Z’ factor = 0.77. Compared to this, our continuous fluorogenic β-galactosidase methodology seems equivalent, considering similarities in the detection limit (∼500 amastigotes/well), convenience (time, sample handling, and cost), and overall performance (Z ≥ 0.8). Importantly, the availability of both orthogonal phenotypic methodologies (differing in *T. cruzi* strains/DTUs, enzyme reporters, and detection techniques) would now permit the counter-screening of compound libraries to confirm pan-strain activity or detect interfering compounds.

Considering the historical success of repurposing antifungal drugs to trypanosomatid infections [33,37–39,61–63], and our own experience with the repositioning of bioactive compounds from yeast to *T. cruzi*, we decided to evaluate six emerging antifungal drugs with the developed fluorogenic methodology. Three of the six evaluated drugs showed measurable activity against *T. cruzi* amastigotes in culture (50% success rate), and two of them, AR-12 and Fosmanogepix, displayed IC_50_ values in the low micromolar range. To the best of our knowledge, this is the first report on the activity of both drugs, alone or in combination, against *T. cruzi* parasites. Due to its moderate solubility in DMSO, Fosmanogepix could not be tested in cell cultures at concentrations above 25 μM. At this concentration, no signs of cytotoxicity were observed. On the other hand, AR-12 showed measurable cytotoxicity on Vero cells, with a selectivity index considered acceptable for initial-stage candidates. In addition, when combined with BNZ at specific concentrations, synergistic and potent antiparasitic effects were observed for both drugs, as well as for the AR-12/Fosmanogepix combination. Therefore, both AR-12 and Fosmanogepix can be considered good candidates to repurpose as initial hits for further development of anti-T. cruzi drugs, especially considering that they have been successfully tested in clinical trials in humans [64].

The observation of these synergistic effects suggests that BNZ, AR-12, and Fosmanogepix act through different mechanisms and/or targets. In fact, benznidazole acts as a pro-drug, activated by a parasite type I nitroreductase, which ultimately produces reactive glyoxal and toxic guanosine-glyoxal adducts [65,66]; while in yeast, AR-12 is an ATP-competitive, time-dependent inhibitor of acetyl coenzyme A synthetase (AceCS) [67], and Fosmanogepix ― also a pro-drug converted *in vivo* into active manogepix/E1210 ― is proposed to inhibit the conserved fungal glycosylphosphatidylinositol (GPI)-anchored cell wall transfer protein 1 (Gwt1), which catalyzes inositol acylation of GPI [68–70]. In addition to fungal AceCS, AR-12 has been reported to competitively inhibit several human kinases, such as 3-phosphoinositide-dependent kinase-1 (PDK1) [71] and p21-activated kinase (PAK) [72], by targeting their ATP-binding sites with affinities in the low micromolar range [73]. Finally, this compound also inhibits human dihydroorotate dehydrogenase (DHODH), displaying a sub-micromolar IC_50_ value in an enzymatic assay [74]. The binding of AR-12 to a hydrophobic pocket in the surface of human DHODH was further confirmed by X-ray crystallography, although the mechanism of enzyme inhibition remains unclear.

Because both drugs have well-established therapeutically relevant targets in fungi or humans, their anti-*T. cruzi* mode(s) of action may be envisaged. In the case of AR-12, AceCS, PDK1, and DHODH have recognizable orthologs in *T. cruzi: Tc*AceCS, *Tc*AEK1, and *Tc*DHODH, respectively. While no studies were found addressing directly the essentiality of *Tc*AceCS, their orthologues *TbAce*CS and *Ld*AceCS seem to be essential for *Trypanosoma brucei* and *Leishmania donovani*, respectively [75–77]. On the contrary, both *Tc*AEK1 and *Tc*DHODH have been previously described as essential targets for the survival of *T. cruzi* parasites [78–80]. Considering this evidence, the hypothesis that AR-12 interferes with *T. cruzi* viability through one (or more) of these targets is favored. Of note, the effectiveness of AR-12 as a host-mediated therapeutic agent against diverse pathogens has also been reported [81]. The proposed mechanisms involve the upregulation of autophagy and the Akt kinase pathway, suggesting that this compound might trigger additional processes activating host Vero cells against *T. cruzi* infection. On the other hand, the target of Fosmanogepix, Gwt1, lacks an evident ortholog in *T. cruzi*. Despite this, the enzymatic activity exists, and it seems clear that the synthesis of GPI-anchored proteins is a critical pathway for parasite physiology [82–84]. In this regard, we favor the hypothesis that Fosmanogepix acts on *T. cruzi* somewhere along this pathway, opening the quest for the responsible target, as this could open new avenues of therapeutic intervention.

Overall, our results indicate that the semi-automatic methodology described here can be conveniently used for the rapid evaluation of up to hundreds of compounds at a time in phenotypic screening against *T. cruzi*, with the potential to be extended to other *T. cruzi* strains or even other trypanosomatid species. The convergence/combination of medium-throughput screening technologies and hypothesis-guided drug repositioning may lead to the discovery of promising novel candidates as well as potentially new exploitable therapeutic targets. Finally, the multitarget compound AR-12 and the inositol acyl-transferase inhibitor Fosmanogepix are good candidates to repurpose for the treatment of Chagas Disease, highlighting the importance of antifungals as a potential source of repositionable drugs to parasitic NTDs with limited funding.

## MATERIAL AND METHODS

### Compounds and reagents

The antifungal drugs Rezafungin acetate (CS-0114091, purity: 98.04%), Fosmanogepix (CS-0079972, purity: ≥ 97.0%), Nikkomycin Z (CS-0015875, purity: ≥ 92.0%), ME1111 (CS-0027148, purity: 99.86%), and T-2307 (CS-0079957, purity: 99.45%) were acquired from ChemScene (https://ChemScene.com). AR-12, a.k.a. OSU-03012 (AH18734, purity: 98.0%) and the fluorogenic substrate 4-methylumbelliferyl-β-D-glucopyranoside (MUG) (AB68029, purity: 98.0%) were from A2B (https://a2bchem.com). Stock solutions of investigational compounds were prepared in DMSO at 20 mM (AR-12), 10 mM (ME1111, Rezafungin acetate, Nikkomycin Z), 5 mM (Fosmanogepix), and 2.5 mM (T-2307) according to the observed solubility. Stock solutions of benznidazole (20 mM in DMSO) and Isopropyl β-D-1-thiogalactopyranoside (IPTG) (1M in distilled water) were used as positive controls in infection and β-galactosidase inhibition experiments, respectively.

### Parasite culture

*T. cruzi* parasites (Tulahuen strain) expressing constitutively *E. coli* β-galactosidase (Tul β-gal) were a kind gift from Fred Buckner (University of Washington, USA) and were used as described [16]. Initial stocks of infective trypomastigotes were produced *in vivo* by serial passage in CF1 mice. At the parasitemia peak, bloodstream trypomastigotes were purified using a Ficoll gradient with a swinging bucket rotor, as previously described [85] and stored in liquid nitrogen until needed.

For all infection experiments, cell-derived Tul β-gal trypomastigotes were cultured by weekly passages in Vero cells at 37°C in a humidified atmosphere containing 5% CO_2_ in minimum essential medium (MEM; Gibco Life Technologies) supplemented with 10% FBS, 10 μg/mL streptomycin and 100 U/mL penicillin. Trypomastigotes were harvested from the supernatants of infected cells at 5,000 × g for 10 min after 96 h post-infection. Cell-derived trypomastigotes were maintained in culture for at least 40 passages without significant loss of β-galactosidase activity. After this, a new culture was initiated on Vero cells from bloodstream trypomastigotes in liquid nitrogen storage.

### Continuous β-galactosidase fluorogenic assay

β-galactosidase activity was assayed fluorometrically with MUG as substrate in PBS supplemented with 2% RPMI, 0.25% DMSO, and 0.5% NP-40, pH 7.4 as activity buffer. A lysate of cell-derived Tul β-gal trypomastigotes (1×10^6^ parasites/mL) was used as a source of the enzyme and MUG concentration was set to 10 μM to avoid inner filtering effect [31]. Assay was performed in solid black polystyrene Corning® NBS 384-well plates (CLS3654-100EA) in a final reaction volume of 100 μL, and the release of 4-methylumbelliferone was monitored continuously during 60 minutes at 37 °C with a FilterMax F5 Multimode Reader (Molecular Devices) using standard 340 nm excitation and 450 nm emission filter set. Optimal β-galactosidase assay concentration (1/2,000) was selected from two-fold serial dilutions (range 1/25 – 1/409,600) to match three criteria: (i) being linearly proportional to V_0_, (ii) display robust signal evolution at 10 μM substrate concentration, and (iii) display linear kinetics for at least 60 minutes. The slopes (dF/dt) from progression curves were determined by linear regression analysis using GraphPad Prism program (version 5.03). Finally, the effect of substrate or additives concentration on the fluorogenic β-galactosidase assay was determined by adding MUG (range: 0 - 10 μM), DMSO (range: 0.5 - 2%), or NP-40 (range: 0.5 - 4%) to the activity buffer and performing the assay as described above. The resultant slopes were used to construct the Michaelis-Menten plot or normalized using the slope of unmodified assay as 100%.

### Simultaneous quantification of β-galactosidase activity in fresh in-situ lysates of Tul β-gal trypomastigotes by continuous and endpoint methodologies

Vero cells (5×10^3^) were seeded on standard 96-well culture plates and incubated O.N. with 100 μL of MEM medium supplemented with 10% FBS. MEM medium was replaced with 100 μL of 4% RPMI medium without phenol red (Gibco #11835030) supplemented with 0.5% DMSO and containing two-fold dilutions of cell-derived Tul β-gal trypomastigotes (range: 0 - 2×10^5^ parasites/well). Then, 100 µL of 1% NP-40 in PBS was added to each well, homogenized by up-and-down pipetting, and incubated for 30 minutes to enable cell lysis. For continuous estimation of β-galactosidase activity, 90 µL of the resultant lysates were transferred to a black 384-well plate and 10 µL of MUG (10 µM final concentration) was added to each well to initiate the reaction. Fluorescence was continuously monitored as indicated in the previous section. For endpoint colorimetric estimation, the chromogenic substrate chlorophenol red-β-D-galactopyranoside (CPRG, Roche #10884308001) was added to each well (50 µM final concentration) and the plate was incubated for 4 hours at 37°C, protected from light. β-galactosidase activity was quantified by measuring absorbance at 595 nm using a FilterMax F5 Multimode Microplate Reader.

### Simultaneous quantification of Tul β-gal amastigotes in infected cells by enzymatic methodologies and qPCR

Vero cells were seeded at a density of 5×10³/well in 96-well plates using MEM medium supplemented with 10% FBS. After 48 hours, cells were infected with 5×10⁴ cell-derived Tul β-gal trypomastigotes per well (multiplicity of infection, MOI=10), in MEM medium supplemented with 4% FBS. Plate was incubated 24 hours for infection and washed twice with PBS to remove non-internalized parasites. Two-fold dilutions of benznidazole (BNZ, range: 0.08 - 20 µM), prepared in 100 µL of RPMI medium without phenol red supplemented with 4% FBS, were added to each well. The final DMSO concentration did not exceed 0.5% in any condition to avoid interference with parasite or host cell viability. Vehicle controls (0.5% DMSO) were included for infected and non-infected wells to establish the maximum and minimum infection levels, respectively. After 72 hours of incubation, 100 µL of 1% NP-40 in PBS was added to each well and treated as described above. After homogenization, the resultant lysates were split for in-parallel evaluation with the three methodologies: 10 µL, 90 µL, and 100 µL for qPCR, fluorogenic (continuous), and colorimetric (endpoint) assays, respectively.

The quantification of β-galactosidase activity was performed as described in the previous section. The slope (dF/dt) or DO_505_ measurement obtained for each BNZ concentration was transformed into percentage of β-galactosidase residual activity, according to Eq.(1) or Eq.(2), respectively:

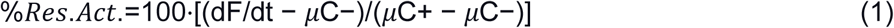

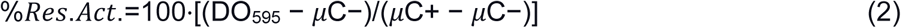

where dF/dt and DO_595_ represent the slope and absorbance values for each BNZ condition, and µC+ and µC− the averages of infected and non-infected vehicle controls, respectively. Half-maximal inhibitory concentration (IC_50_) and Hill slope parameters were estimated for each assay by fitting the four-parameter Hill equation to experimental data from the resultant dose-response curves *(%Res.Act.* vs. log[BNZ]) using GraphPad Prism.

### Real-time PCR procedures

#### A) DNA extraction

For qPCR quantification of *T. cruzi* DNA, 10 µL of each amastigote lysate was diluted to 500 µL with PBS and immediately mixed with an equal volume of GE buffer (6 mol/L guanidine hydrochloride, 0.2 mol/L EDTA, pH 8.00). Samples were incubated for 24 hours at room temperature and stored at 4°C until DNA extraction. Three hundred µL of sample were processed using the QIAamp DNA Blood Mini Kit (Qiagen, Germany) as described in Ramirez and Moreira [86]. The DNA eluates were stored at −20°C until qPCR analysis.

#### B) Standard curve

To build the standard curve, 500 µL of PBS containing 5×10^5^ Tul β-gal trypomastigotes was mixed with an equal volume of GE buffer and processed in the same way as described above for experimental samples.

#### C) Duplex real-time PCR

A duplex qPCR assay targeting *T. cruzi* satellite DNA (SatDNA) and the sequence of an internal amplification control (IAC) was performed with 5 μL of purified DNA, using the FastStart Universal Probe Master Mix (Roche Diagnostics GmbHCorp., Germany) in a final volume of 20 μL, as previously described [86]. The qPCR reactions were carried out in a CFX96 real-time PCR detection system (Bio-Rad, CA). Standard curve was plotted with ten-fold serial dilutions of total DNA obtained from the spiked sample (10^6^ parasite equivalents/mL) and used to quantify parasitic loads in experimental samples.

### Evaluation of anti-T. cruzi activity of antifungal drugs using the continuous fluorogenic methodology

Dose-response curves for antifungal drugs were built with at least eight two-fold serially diluted concentrations. The concentration range varied according to drug solubility, to limit DMSO concentration in culture to 0.5%. Drugs were added to Vero cells infected with Tul β-gal trypomastigotes (MOI=10) as described above. After 72 hours of drug exposure, cultures were lysed and analyzed for β-galactosidase activity by using the continuous fluorometric assay, as described in the previous sections. Infected and non-infected vehicle controls were used to normalize data. After transforming slope values into *%Res.Act.* with Eq.(1) and constructing dose-response plots, IC_50_ values were estimated by using GraphPad Prims. BNZ was used as positive control.

### Cytotoxicity of antifungal drugs on Vero cells

A resazurin (RZ) assay [87] was used to evaluate the cytotoxicity of investigational compounds on non-infected Vero cells. Culture procedures, drug concentrations, and drug-exposure times were identical to those used in the evaluation of anti-*T. cruzi* activity. A vehicle control (0.5% DMSO) was included to establish the maximum level of resazurin reduction (100 % viability). After drug exposure, 10 µL of freshly prepared 10X (440 µM in PBS) RZ solution was added to each well and plates were incubated for 7 hours at 37°C, protected from light. Finally, samples were transferred to a black 96-well plate and fluorescence (λexcitation/emission: 535nm/595nm) was measured in a FilterMax F5 Multimode Microplate Reader. Fluorescence measurements for each condition were transformed into *%Cell.Viability* by using Eq.3:

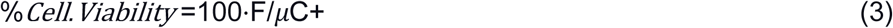

Where F represents Fluorescence measurements and μC+ the average of vehicle controls, respectively. Half maximal cytotoxic concentration (CC_50_) was estimated from constructed dose-response plots as previously described. The selectivity index (SI) was calculated for each compound as the ratio of CC_50_ to IC_50_.

### Inhibition of β-galactosidase by antifungal drugs

Antifungal compounds were tested in a dose-response manner (range: variable) using the continuous β-galactosidase assay described in the previous sections. Compounds were initially pre-incubated at 37°C with the lysate containing β-galactosidase (dilution 1/2,000) in the activity buffer for 30 minutes, prior to the addition of MUG substrate (10 μM). Data collection and processing were performed exactly as described above. Percentage of β-galactosidase residual activity was calculated for each condition according to Eq.(1) to derive dose-response plots. IPTG (range: 0 - 5 mM) was used as β-galactosidase inhibition control.

### Partial automation of the continuous methodology

The workflow described in this paper employs an OT-2 liquid-handling robot (Opentrons, NY). Scripts were developed in Python to operate the OT-2 robot to automate specific protocol steps. Protocols were designed to prepare drug serial dilutions or drug combinations, dispense the lysis solution to treated cell cultures, transfer the lysate to a black 384-well plate, and add MUG substrate to initiate the enzymatic reaction. All the scripts and their descriptions are available at https://github.com/trypanosomatics/OT2-protocols-for-antifungal-essays.

### Semi-automatic evaluation of drug combinations

Active compounds AR-12, Fosmanogepix, and BNZ were tested in drug-drug combination assays. Drug combinations were assembled by the OT-2 robot in PCR plates by mixing three-fold serial dilutions of drug 1 on the columns and three-fold serial dilutions of drug 2 on the rows (Supplementary Figure S7). In all cases, dilution steps (and therefore drug combinations) were prepared in DMSO. Once the combination plates were set, drug mixes were added to infected Vero cells to achieve the desired final concentrations of AR-12 (0 – 50 µM), Fosmanogepix (0 – 12.5 µM), and BNZ (0 – 5 µM) in the culture wells. Final DMSO concentration was maintained at 0.5% to prevent any impact on cell or parasite viability. After 72 hours of exposure, OT2 added 100 µL of lysis buffer to the plates and mixed each well by up-and down pipetting. After a 30 minutes incubation, the robot transferred 90 µL of the lysates to black 384-well reading plates and added 10 µL of MUG substrate. Data acquisition and processing were performed as previously described to generate effect matrices containing the observed antiparasitic effect (100 - *%Res.Act.*) for each combination of doses. The resultant effect matrices were analyzed with SinergyFinder [44,45] and SynergyFinder Plus [46] with default parameters to generate synergy distribution maps. Finally, cytotoxicity of drug combinations on Vero cells was determined as described above for individual compounds.

## Supporting information

Electronic Supplementary Material

## Abbreviations

BNZ: Benznidazole
CV: coefficient of variation (also known as RSD = relative standard deviation);

## ACKNOWLEDGEMENTS

Mercedes Didier Garnham is supported by a fellowship from CONICET, the National Research Council of Argentina. Juan Carlos Ramírez, Fernán Agüero, and Emir Salas Sarduy are Career Scientists at CONICET.

## REFERENCES

1. Bonney KM. Chagas disease in the 21st Century: a public health success or an emerging threat? Parasite. 2014;21: 11. doi:10.1051/parasite/2014012

2. Pérez-Molina JA, Molina I. Chagas disease. Lancet Lond Engl. 2018;391: 82–94. doi:10.1016/S0140-6736(17)31612-4

3. Molina I, Salvador F, Sánchez-Montalvá A, Treviño B, Serre N, Sao Avilés A, et al. Toxic Profile of Benznidazole in Patients with Chronic Chagas Disease: Risk Factors and Comparison of the Product from Two Different Manufacturers. Antimicrob Agents Chemother. 2015;59: 6125–6131. doi:10.1128/AAC.04660-14

4. Hasslocher-Moreno AM, do Brasil PEAA, de Sousa AS, Xavier SS, Chambela MC, Sperandio da Silva GM. Safety of benznidazole use in the treatment of chronic Chagas’ disease. J Antimicrob Chemother. 2012;67: 1261–1266. doi:10.1093/jac/dks027

5. Sales Junior PA, Molina I, Fonseca Murta SM, Sánchez-Montalvá A, Salvador F, Corrêa-Oliveira R, et al. Experimental and Clinical Treatment of Chagas Disease: A Review. Am J Trop Med Hyg. 2017;97: 1289–1303. doi:10.4269/ajtmh.16-0761

6. Andrews KT, Fisher G, Skinner-Adams TS. Drug repurposing and human parasitic protozoan diseases. Int J Parasitol Drugs Drug Resist. 2014;4: 95–111. doi:10.1016/j.ijpddr.2014.02.002

7. Alberca LN, Chuguransky SR, Álvarez CL, Talevi A, Salas-Sarduy E. In silico Guided Drug Repurposing: Discovery of New Competitive and Non-competitive Inhibitors of Falcipain-2. Front Chem. 2019;7: 534. doi:10.3389/fchem.2019.00534

8. Brown D. Unfinished business: target-based drug discovery. Drug Discov Today. 2007;12: 1007–1012. doi:10.1016/j.drudis.2007.10.017

9. Urán Landaburu L, Berenstein AJ, Videla S, Maru P, Shanmugam D, Chernomoretz A, et al. TDR Targets 6: driving drug discovery for human pathogens through intensive chemogenomic data integration. Nucleic Acids Res. 2020;48: D992– D1005. doi:10.1093/nar/gkz999

10. Urán Landaburu L, Didier Garnham M, Agüero F. Targeting trypanosomes: how chemogenomics and artificial intelligence can guide drug discovery. Biochem Soc Trans. 2023;51: 195–206. doi:10.1042/BST20220618

11. Gilbert IH. Drug discovery for neglected diseases: molecular target-based and phenotypic approaches. J Med Chem. 2013;56: 7719–7726. doi:10.1021/jm400362b

12. Alonso-Padilla J, Rodríguez A. High throughput screening for anti-Trypanosoma cruzi drug discovery. PLoS Negl Trop Dis. 2014;8: e3259. doi:10.1371/journal.pntd.0003259

13. Peña I, Pilar Manzano M, Cantizani J, Kessler A, Alonso-Padilla J, Bardera AI, et al. New compound sets identified from high throughput phenotypic screening against three kinetoplastid parasites: an open resource. Sci Rep. 2015;5: 8771. doi:10.1038/srep08771

14. Salas-Sarduy E, Landaburu LU, Carmona AK, Cazzulo JJ, Agüero F, Alvarez VE, et al. Potent and selective inhibitors for M32 metallocarboxypeptidases identified from high-throughput screening of anti-kinetoplastid chemical boxes. PLoS Negl Trop Dis. 2019;13: e0007560. doi:10.1371/journal.pntd.0007560

15. Dantas RF, Torres-Santos EC, Silva FP. Past and future of trypanosomatids high-throughput phenotypic screening. Mem Inst Oswaldo Cruz. 2022;117. doi:10.1590/0074-02760210402

16. Buckner FS, Verlinde CL, La Flamme AC, Van Voorhis WC. Efficient technique for screening drugs for activity against Trypanosoma cruzi using parasites expressing beta-galactosidase. Antimicrob Agents Chemother. 1996;40: 2592–7. Available: http://www.ncbi.nlm.nih.gov/pubmed/8913471 http://www.pubmedcentral.nih.gov/articlerender.fcgi?artid=PMC163582

17. MacLean LM, Thomas J, Lewis MD, Cotillo I, Gray DW, De Rycker M. Development of Trypanosoma cruzi in vitro assays to identify compounds suitable for progression in Chagas’ disease drug discovery. Buscaglia CA, editor. PLoS Negl Trop Dis. 2018;12: e0006612. doi:10.1371/journal.pntd.0006612

18. Eustice DC, Feldman PA, Colberg-Poley AM, Buckery RM, Neubauer RH. A sensitive method for the detection of beta-galactosidase in transfected mammalian cells. BioTechniques. 1991;11: 739–740, 742–743.

19. Cantizani J, Gamallo P, Cotillo I, Alvarez-Velilla R, Martin J. Rate-of-Kill (RoK) assays to triage large compound sets for Chagas disease drug discovery: Application to GSK Chagas Box. PLoS Negl Trop Dis. 2021;15: e0009602. doi:10.1371/journal.pntd.0009602

20. Didier Garnham M, Landaburu LU, Salas-Sarduy E, Agüero F. New hit compounds against Trypanosoma cruzi derived from yeast chemogenomic profiling. 2025;[Manuscript in preparation]: [Manuscript in preparation].

21. Sharma SK, Poudel Sharma S, Leblanc RM. Methods of detection of β-galactosidase enzyme in living cells. Enzyme Microb Technol. 2021;150: 109885. doi:10.1016/j.enzmictec.2021.109885

22. Chiu NHL, Watson AL. Measuring β-Galactosidase Activity in Gram-Positive Bacteria Using a Whole-Cell Assay with MUG as a Fluorescent Reporter. Curr Protoc Toxicol. 2017;74: 4.44.1–4.44.8. doi:10.1002/cptx.35

23. Jones I, Hamilton J, Srivastava R, Galloway P. Effect of neutral and acid pH on the fluorescence of 4-methylumbelliferone and the implications for dry blood spot assays. Mol Genet Metab. 2013;108: S51. doi:10.1016/j.ymgme.2012.11.120

24. Zhi H, Wang J, Wang S, Wei Y. Fluorescent Properties of Hymecromone and Fluorimetric Analysis of Hymecromone in Compound Dantong Capsule. J Spectrosc. 2013;2013: 1–9. doi:10.1155/2013/147128

25. Woollen JW, Walker PG. The fluorimetric estimation of N-acetyl-beta-glucosaminidase and beta-galactosidase in blood plasma. Clin Chim Acta Int J Clin Chem. 1965;12: 647–658. doi:10.1016/0009-8981(65)90147-6

26. Robins E, Hirsch HE, Emmons SS. Glycosidases in the nervous system. I. Assay, some properties, and distribution of beta-galactosidase, beta-glucoronidase, and beta-glucosidase. J Biol Chem. 1968;243: 4246–4252.

27. Cheetham PS, Dance NE. The separation and characterization of the methylumbelliferyl beta-galactosidases of human liver. Biochem J. 1976;157: 189–195. doi:10.1042/bj1570189

28. Labrousse H, Guesdon JL, Ragimbeau J, Avrameas S. Miniaturization of beta-galactosidase immunoassays using chromogenic and fluorogenic substrates. J Immunol Methods. 1982;48: 133–147. doi:10.1016/0022-1759(82)90188-0

29. Gary RK, Kindell SM. Quantitative assay of senescence-associated beta-galactosidase activity in mammalian cell extracts. Anal Biochem. 2005;343: 329–334. doi:10.1016/j.ab.2005.06.003

30. Hosseini A, Mas J. The β-galactosidase assay in perspective: Critical thoughts for biosensor development. Anal Biochem. 2021;635: 114446. doi:10.1016/j.ab.2021.114446

31. Palmier MO, Van Doren SR. Rapid determination of enzyme kinetics from fluorescence: overcoming the inner filter effect. Anal Biochem. 2007;371: 43–51. doi:10.1016/j.ab.2007.07.008

32. Ramírez JC, Cura CI, Da Cruz Moreira O, Lages-Silva E, Juiz N, Velázquez E, et al. Analytical Validation of Quantitative Real-Time PCR Methods for Quantification of Trypanosoma cruzi DNA in Blood Samples from Chagas Disease Patients. J Mol Diagn. 2015;17: 605–615. doi:10.1016/j.jmoldx.2015.04.010

33. Cosentino RO, Agüero F. Genetic profiling of the isoprenoid and sterol biosynthesis pathway genes of Trypanosoma cruzi. PloS One. 2014;9: e96762. doi:10.1371/journal.pone.0096762

34. Tewary P, Veena K, Pucadyil TJ, Chattopadhyay A, Madhubala R. The sterol-binding antibiotic nystatin inhibits entry of non-opsonized Leishmania donovani into macrophages. Biochem Biophys Res Commun. 2006;339: 661–666. doi:10.1016/j.bbrc.2005.11.062

35. Machado PRL, Rosa MEA, Guimarães LH, Prates FVO, Queiroz A, Schriefer A, et al. Treatment of Disseminated Leishmaniasis With Liposomal Amphotericin B. Clin Infect Dis Off Publ Infect Dis Soc Am. 2015;61: 945–949. doi:10.1093/cid/civ416

36. Kaiser M, Mäser P, Tadoori LP, Ioset J-R, Brun R. Antiprotozoal Activity Profiling of Approved Drugs: A Starting Point toward Drug Repositioning. PloS One. 2015;10: e0135556. doi:10.1371/journal.pone.0135556

37. Yamamoto ES, Jesus JA, Bezerra-Souza A, Laurenti MD, Ribeiro SP, Passero LFD. Activity of Fenticonazole, Tioconazole and Nystatin on New World Leishmania Species. Curr Top Med Chem. 2019;18: 2338–2346. doi:10.2174/1568026619666181220114627

38. Porta EOJ, Kalesh K, Steel PG. Navigating drug repurposing for Chagas disease: advances, challenges, and opportunities. Front Pharmacol. 2023;14: 1233253. doi:10.3389/fphar.2023.1233253

39. Maciel BJ, Reigada C, Digirolamo FA, Rengifo M, Pereira CA, Miranda MR, et al. The potential of the antifungal nystatin to be repurposed to fight the protozoan Trypanosoma cruzi. Front Microbiol. 2025;16: 1539629. doi:10.3389/fmicb.2025.1539629

40. Molina I, Gómez I Prat J, Salvador F, Treviño B, Sulleiro E, Serre N, et al. Randomized Trial of Posaconazole and Benznidazole for Chronic Chagas’ Disease. N Engl J Med. 2014;370: 1899–1908. doi:10.1056/NEJMoa1313122

41. Torrico F, Gascon J, Ortiz L, Alonso-Vega C, Pinazo M-J, Schijman A, et al. Treatment of adult chronic indeterminate Chagas disease with benznidazole and three E1224 dosing regimens: a proof-of-concept, randomised, placebo-controlled trial. Lancet Infect Dis. 2018;18: 419–430. doi:10.1016/S1473-3099(17)30538-8

42. Riss TL, Moravec RA, Niles AL, Duellman S, Benink HA, Worzella TJ, et al. Cell Viability Assays. In: Markossian S, Grossman A, Baskir H, Arkin M, Auld D, Austin C, et al., editors. Assay Guidance Manual. Bethesda (MD): Eli Lilly & Company and the National Center for Advancing Translational Sciences; 2004. Available: http://www.ncbi.nlm.nih.gov/books/NBK144065/

43. Hoang KV, Borteh HM, Rajaram MVS, Peine KJ, Curry H, Collier MA, et al. Acetalated dextran encapsulated AR-12 as a host-directed therapy to control Salmonella infection. Int J Pharm. 2014;477: 334–343. doi:10.1016/j.ijpharm.2014.10.022

44. Ianevski A, Giri AK, Aittokallio T. SynergyFinder 2.0: visual analytics of multi-drug combination synergies. Nucleic Acids Res. 2020;48: W488–W493. doi:10.1093/nar/gkaa216

45. Ianevski A, Giri AK, Aittokallio T. SynergyFinder 3.0: an interactive analysis and consensus interpretation of multi-drug synergies across multiple samples. Nucleic Acids Res. 2022;50: W739–W743. doi:10.1093/nar/gkac382

46. Zheng S, Wang W, Aldahdooh J, Malyutina A, Shadbahr T, Tanoli Z, et al. SynergyFinder Plus: Toward Better Interpretation and Annotation of Drug Combination Screening Datasets. Genomics Proteomics Bioinformatics. 2022;20: 587–596. doi:10.1016/j.gpb.2022.01.004

47. Gabaldón-Figueira JC, Martinez-Peinado N, Escabia E, Ros-Lucas A, Chatelain E, Scandale I, et al. State-of-the-Art in the Drug Discovery Pathway for Chagas Disease: A Framework for Drug Development and Target Validation. Res Rep Trop Med. 2023;Volume 14: 1–19. doi:10.2147/RRTM.S415273

48. De Rycker M, Wyllie S, Horn D, Read KD, Gilbert IH. Anti-trypanosomatid drug discovery: progress and challenges. Nat Rev Microbiol. 2023;21: 35–50. doi:10.1038/s41579-022-00777-y

49. Copeland RA. Mechanistic considerations in high-throughput screening. Anal Biochem. 2003;320: 1–12. doi:10.1016/S0003-2697(03)00346-4

50. Fior S, Vianelli A, Gerola PD. A novel method for fluorometric continuous measurement of β-glucuronidase (GUS) activity using 4-methyl-umbelliferyl-β-d-glucuronide (MUG) as substrate. Plant Sci. 2009;176: 130–135. doi:10.1016/j.plantsci.2008.10.001

51. Jones I, Hamilton J, Srivastava R, Galloway P. Effect of neutral and acid pH on the fluorescence of 4-methylumbelliferone and the implications for dry blood spot assays. Mol Genet Metab. 2013;108: S51. doi:10.1016/j.ymgme.2012.11.120

52. Gulin JEN, Rocco DM, Alonso V, Cribb P, Altcheh J, García-Bournissen F. Optimization and biological validation of an *in vitro* assay using the transfected Dm28c/pLacZ *Trypanosoma cruzi* strain. Biol Methods Protoc. 2021;6: bpab004. doi:10.1093/biomethods/bpab004

53. Alonso VL, Manarin R, Perdomo V, Gulin E, Serra E, Cribb P. In Vitro Drug Screening Against All Life Cycle Stages of Trypanosoma cruzi Using Parasites Expressing &#946;-galactosidase. J Vis Exp. 2021; 63210. doi:10.3791/63210

54. Okuno T, Goto Y, Matsumoto Y, Otsuka H, Matsumoto Y. Applications of recombinant Leishmania amazonensis expressing egfp or the beta-galactosidase gene for drug screening and histopathological analysis. Exp Anim. 2003;52: 109–118. doi:10.1538/expanim.52.109

55. Da Silva Santos AC, Moura DMN, Dos Santos TAR, De Melo Neto OP, Pereira VRA. Assessment of Leishmania cell lines expressing high levels of beta-galactosidase as alternative tools for the evaluation of anti-leishmanial drug activity. J Microbiol Methods. 2019;166: 105732. doi:10.1016/j.mimet.2019.105732

56. Bot C, Hall BS, Bashir N, Taylor MC, Helsby NA, Wilkinson SR. Trypanocidal activity of aziridinyl nitrobenzamide prodrugs. Antimicrob Agents Chemother. 2010;54: 4246–4252. doi:10.1128/AAC.00800-10

57. Lara LS, Moreira CS, Calvet CM, Lechuga GC, Souza RS, Bourguignon SC, et al. Efficacy of 2-hydroxy-3-phenylsulfanylmethyl-[1,4]-naphthoquinone derivatives against different Trypanosoma cruzi discrete type units: Identification of a promising hit compound. Eur J Med Chem. 2018;144: 572–581. doi:10.1016/j.ejmech.2017.12.052

58. Tayama Y, Mizukami S, Toume K, Komatsu K, Yanagi T, Nara T, et al. Anti-Trypanosoma cruzi activity of Coptis rhizome extract and its constituents. Trop Med Health. 2023;51: 12. doi:10.1186/s41182-023-00502-2

59. Quiroga C, Incerti M, Benitez D, Manta E, Medeiros A, Comini MA. Development of bioluminescent reporter Trypanosoma cruzi and bioassay for compound screening. Front Chem Biol. 2024;3. doi:10.3389/fchbi.2024.1423430

60. Branchini BR, Ablamsky DM, Davis AL, Southworth TL, Butler B, Fan F, et al. Red-emitting luciferases for bioluminescence reporter and imaging applications. Anal Biochem. 2010;396: 290–297. doi:10.1016/j.ab.2009.09.009

61. Kumari S, Kumar V, Tiwari RK, Ravidas V, Pandey K, Kumar A. Amphotericin B: A drug of choice for Visceral Leishmaniasis. Acta Trop. 2022;235: 106661. doi:10.1016/j.actatropica.2022.106661

62. Kaiser M, Mäser P, Tadoori LP, Ioset J-R, Brun R. Antiprotozoal Activity Profiling of Approved Drugs: A Starting Point toward Drug Repositioning. Sullivan DJ, editor. PLOS ONE. 2015;10: e0135556. doi:10.1371/journal.pone.0135556

63. Tewary P, Veena K, Pucadyil TJ, Chattopadhyay A, Madhubala R. The sterol-binding antibiotic nystatin inhibits entry of non-opsonized Leishmania donovani into macrophages. Biochem Biophys Res Commun. 2006;339: 661–666. doi:10.1016/j.bbrc.2005.11.062

64. Rauseo AM, Coler-Reilly A, Larson L, Spec A. Hope on the Horizon: Novel Fungal Treatments in Development. Open Forum Infect Dis. 2020;7: ofaa016. doi:10.1093/ofid/ofaa016

65. Hall BS, Wilkinson SR. Activation of Benznidazole by Trypanosomal Type I Nitroreductases Results in Glyoxal Formation. Antimicrob Agents Chemother. 2012;56: 115–123. doi:10.1128/AAC.05135-11

66. Rajão MA, Furtado C, Alves CL, Passos-Silva DG, de Moura MB, Schamber-Reis BL, et al. Unveiling benznidazole’s mechanism of action through overexpression of DNA repair proteins in Trypanosoma cruzi. Environ Mol Mutagen. 2014;55: 309–321. doi:10.1002/em.21839

67. Koselny K, Green J, Favazzo L, Glazier VE, DiDone L, Ransford S, et al. Antitumor/Antifungal Celecoxib Derivative AR-12 is a Non-Nucleoside Inhibitor of the ANL-Family Adenylating Enzyme Acetyl CoA Synthetase. ACS Infect Dis. 2016;2: 268–280. doi:10.1021/acsinfecdis.5b00134

68. Miyazaki M, Horii T, Hata K, Watanabe N-A, Nakamoto K, Tanaka K, et al. In vitro activity of E1210, a novel antifungal, against clinically important yeasts and molds. Antimicrob Agents Chemother. 2011;55: 4652–4658. doi:10.1128/AAC.00291-11

69. Watanabe N-A, Miyazaki M, Horii T, Sagane K, Tsukahara K, Hata K. E 1210, a new broad-spectrum antifungal, suppresses Candida albicans hyphal growth through inhibition of glycosylphosphatidylinositol biosynthesis. Antimicrob Agents Chemother. 2012;56: 960–971. doi:10.1128/AAC.00731-11

70. Pappas PG, Vazquez JA, Oren I, Rahav G, Aoun M, Bulpa P, et al. Clinical safety and efficacy of novel antifungal, fosmanogepix, for the treatment of candidaemia: results from a Phase 2 trial. J Antimicrob Chemother. 2023;78: 2471–2480. doi:10.1093/jac/dkad256

71. Lee TX, Packer MD, Huang J, Akhmametyeva EM, Kulp SK, Chen C-S, et al. Growth inhibitory and anti-tumour activities of OSU-03012, a novel PDK-1 inhibitor, on vestibular schwannoma and malignant schwannoma cells. Eur J Cancer Oxf Engl 1990. 2009;45: 1709–1720. doi:10.1016/j.ejca.2009.03.013

72. Porchia LM, Guerra M, Wang Y-C, Zhang Y, Espinosa AV, Shinohara M, et al. 2-amino-N-{4-[5-(2-phenanthrenyl)-3-(trifluoromethyl)-1H-pyrazol-1-yl]-phenyl} acetamide (OSU-03012), a celecoxib derivative, directly targets p21-activated kinase. Mol Pharmacol. 2007;72: 1124–1131. doi:10.1124/mol.107.037556

73. Zhu J, Huang J-W, Tseng P-H, Yang Y-T, Fowble J, Shiau C-W, et al. From the cyclooxygenase-2 inhibitor celecoxib to a novel class of 3-phosphoinositide-dependent protein kinase-1 inhibitors. Cancer Res. 2004;64: 4309–4318. doi:10.1158/0008-5472.CAN-03-4063

74. Abt ER, Rosser EW, Durst MA, Lok V, Poddar S, Le TM, et al. Metabolic Modifier Screen Reveals Secondary Targets of Protein Kinase Inhibitors within Nucleotide Metabolism. Cell Chem Biol. 2020;27: 197–205.e6. doi:10.1016/j.chembiol.2019.10.012

75. Rivière L, Moreau P, Allmann S, Hahn M, Biran M, Plazolles N, et al. Acetate produced in the mitochondrion is the essential precursor for lipid biosynthesis in procyclic trypanosomes. Proc Natl Acad Sci U S A. 2009;106: 12694–12699. doi:10.1073/pnas.0903355106

76. Mazet M, Morand P, Biran M, Bouyssou G, Courtois P, Daulouède S, et al. Revisiting the central metabolism of the bloodstream forms of Trypanosoma brucei: production of acetate in the mitochondrion is essential for parasite viability. PLoS Negl Trop Dis. 2013;7: e2587. doi:10.1371/journal.pntd.0002587

77. Soumya N, Panara MN, Neerupudi KB, Singh S. Functional analysis of an AMP forming acetyl CoA synthetase from Leishmania donovani by gene overexpression and targeted gene disruption approaches. Parasitol Int. 2017;66: 992–1002. doi:10.1016/j.parint.2016.11.001

78. Annoura T, Nara T, Makiuchi T, Hashimoto T, Aoki T. The origin of dihydroorotate dehydrogenase genes of kinetoplastids, with special reference to their biological significance and adaptation to anaerobic, parasitic conditions. J Mol Evol. 2005;60: 113–127. doi:10.1007/s00239-004-0078-8

79. Beltran-Hortelano I, Alcolea V, Font M, Pérez-Silanes S. Examination of multiple Trypanosoma cruzi targets in a new drug discovery approach for Chagas disease. Bioorg Med Chem. 2022;58: 116577. doi:10.1016/j.bmc.2021.116577

80. Chiurillo MA, Jensen BC, Docampo R. Drug Target Validation of the Protein Kinase AEK1, Essential for Proliferation, Host Cell Invasion, and Intracellular Replication of the Human Pathogen Trypanosoma cruzi. Microbiol Spectr. 2021;9: e0073821. doi:10.1128/Spectrum.00738-21

81. Collier MA, Peine KJ, Gautam S, Oghumu S, Varikuti S, Borteh H, et al. Host-mediated Leishmania donovani treatment using AR-12 encapsulated in acetalated dextran microparticles. Int J Pharm. 2016;499: 186–194. doi:10.1016/j.ijpharm.2016.01.004

82. N G, Rl T, K M-W. Proteins with glycosylphosphatidylinositol (GPI) signal sequences have divergent fates during a GPI deficiency. GPIs are essential for nuclear division in Trypanosoma cruzi. J Biol Chem. 1997;272. doi:10.1074/jbc.272.19.12482

83. Cardoso MS, Junqueira C, Trigueiro RC, Shams-Eldin H, Macedo CS, Araújo PR, et al. Identification and functional analysis of Trypanosoma cruzi genes that encode proteins of the glycosylphosphatidylinositol biosynthetic pathway. PLoS Negl Trop Dis. 2013;7: e2369. doi:10.1371/journal.pntd.0002369

84. Morotti ALM, Martins-Teixeira MB, Carvalho I. Protozoan Parasites Glycosylphosphatidylinositol Anchors: Structures, Functions and Trends for Drug Discovery. Curr Med Chem. 2019;26: 4301–4322. doi:10.2174/0929867324666170727110801

85. Tekiel VS, Mirkin GA, Gonzalez Cappa SM. Chagas’ disease: reactivity against homologous tissues induced by different strains of *Trypanosoma cruzi*. Parasitology. 1997;115: 495–502. doi:10.1017/S0031182097001625

86. Ramirez JC, Moreira OC. Real-time PCR for Trypanosoma cruzi infection diagnosis and treatment response monitoring of patients with Chagas disease. 2nd Ed. T cruzi infection: Methods and Protocols. 2nd Ed. Humana Press [IN PRESS]; Available: https://link.springer.com/book/9781071648476

87. Ansar Ahmed S, Gogal RM, Walsh JE. A new rapid and simple non-radioactive assay to monitor and determine the proliferation of lymphocytes: an alternative to [3H]thymidine incorporation assay. J Immunol Methods. 1994;170: 211–224. doi:10.1016/0022-1759(94)90396-4

