## Supplementary material for "Identification of antifungal agents AR-12 and Fosmanogepix as anti-*Trypanosoma cruzi* drugs through an enhanced fluorogenic β-galactosidase phenotypic screening assay": Electronic Supplementary Material

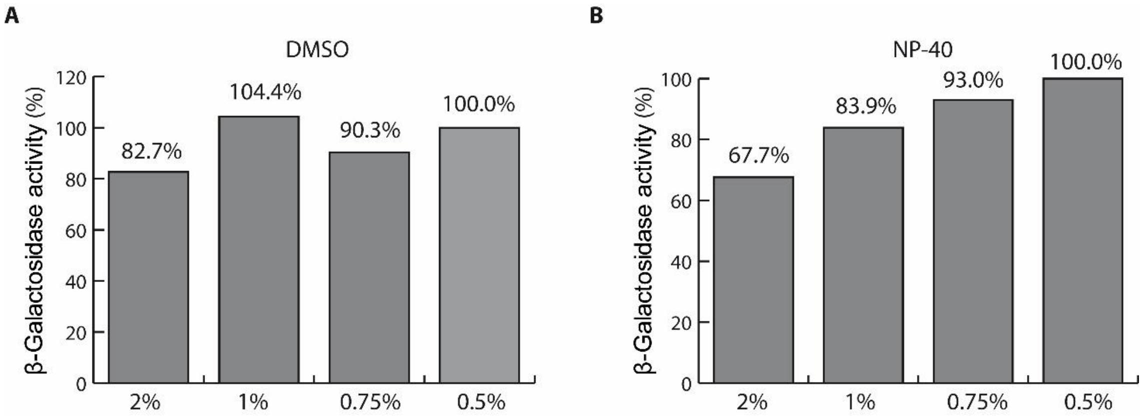


**Supplementary Figure S1. Effect of different additives on the hydrolysis of MUG by β-galactosidase.** The assay was carried out on PBS supplemented with 2% RPMI, 0.25% DMSO, and 0.5% NP-40, pH 7.4, as reaction buffer to match the resultant matrix from culture hydrolysis. In all cases, a lysate of Tul β-gal trypomastigotes (dilution 1:2,000) was used as β-galactosidase source, and the concentration of MUG substrate was fixed at 10 µM.

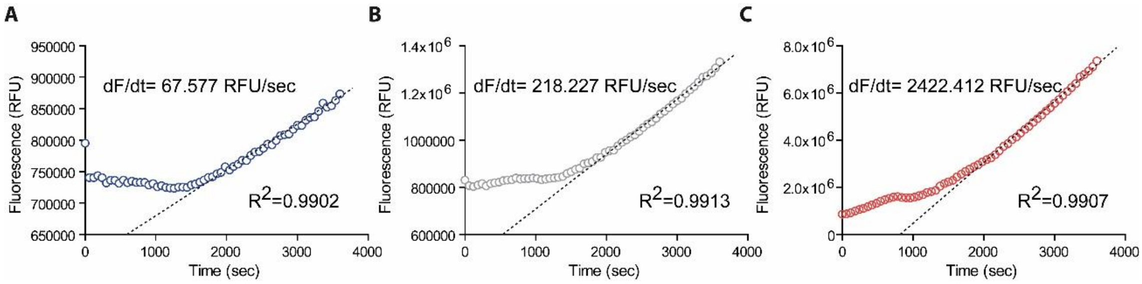


**Supplementary Figure S2. Representative progression curves for the hydrolysis of MUG by β-galactosidase derived from fresh Tul β-gal trypomastigotes.** Tul β-gal trypomastigotes were spiked into wells containing $5x10^3$ Vero cells. After lysis, 100 µL of lysate was transferred to a black 384-well plate, and MUG (10 µM) was added to initiate the reaction. A) Progression curve corresponding to 1,560 trypomastigotes/well. B) 3,125 trypomastigotes/well. C) 50,000 trypomastigotes/well. Steady-state slopes were obtained after 25 minutes (1,500 sec), independently of the number of trypomastigotes added to the well. Strong linear correlations were observed from this time-point.

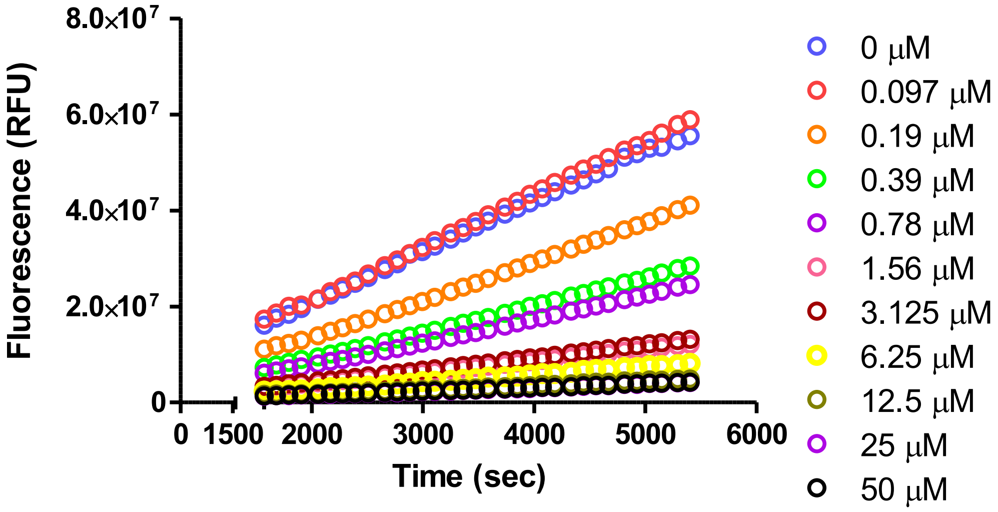


**Supplementary Figure S3. Extended linear progression curves for the hydrolysis of MUG by β-galactosidase derived from Tul β-gal amastigotes.** Infected Vero cells were treated with the indicated BNZ concentrations for 72 hours before culture lysis. After the transference of lysates (90 µL) to the reading plate, MUG (10 µM) was added to initiate the reaction. A "blind" incubation of 25 minutes was included to facilitate linear regression analysis. Independent of the β-galactosidase level, steady-state slopes remained constant for at least 90 minutes (5,400 sec) after the addition of MUG substrate.

**Supplementary Table SI. Summary of fitting parameters from benznidazole dose-response curves.**

| Parameter | CPRG_1 | CPRG_2 | CPRG_3 | Average ± SD | MUG_1 | MUG_2 | MUG_3 | Average ± SD |
|---|---|---|---|---|---|---|---|---|
| Best-fit values | | | | | | | | |
| $IC_{50}$ (µmol/L) | 0.9149 | 0.8192 | 0.6533 | **0.7958 ± 0.1324** | 0.5552 | 0.7659 | 0.9401 | **0.7537 ± 0.1927** |
| HillSlope | -1.139 | -0.8923 | -0.8239 | - | -1.628 | -1.371 | -0.9521 | - |
| 95% Confidence Intervals | | | | | | | | |
| $IC_{50}$ (µmol/L) | 0.6889 to 1.215 | 0.6820 to 0.9839 | 0.5513 to 0.7742 | - | 0.5088 to 0.6058 | 0.6546 to 0.8961 | 0.6334 to 1.395 | - |
| HillSlope | -1.474 to -0.8044 | -1.042 to -0.7431 | -0.9458 to -0.7020 | - | -1.834 to -1.423 | -1.636 to -1.106 | -1.308 to -0.5961 | - |
| Goodness of Fit | | | | | | | | |
| $R^2$ | 0.971 | 0.987 | 0.9885 | - | 0.9969 | 0.9896 | 0.9424 | - |

Data corresponding to the end-point chromogenic (CPRG) and the continuous fluorogenic (MUG) methodologies are indicated in light-blue and light-green colors, respectively. The average $IC_{50}$ values from three independent experiments for each methodology are indicated in bold text.

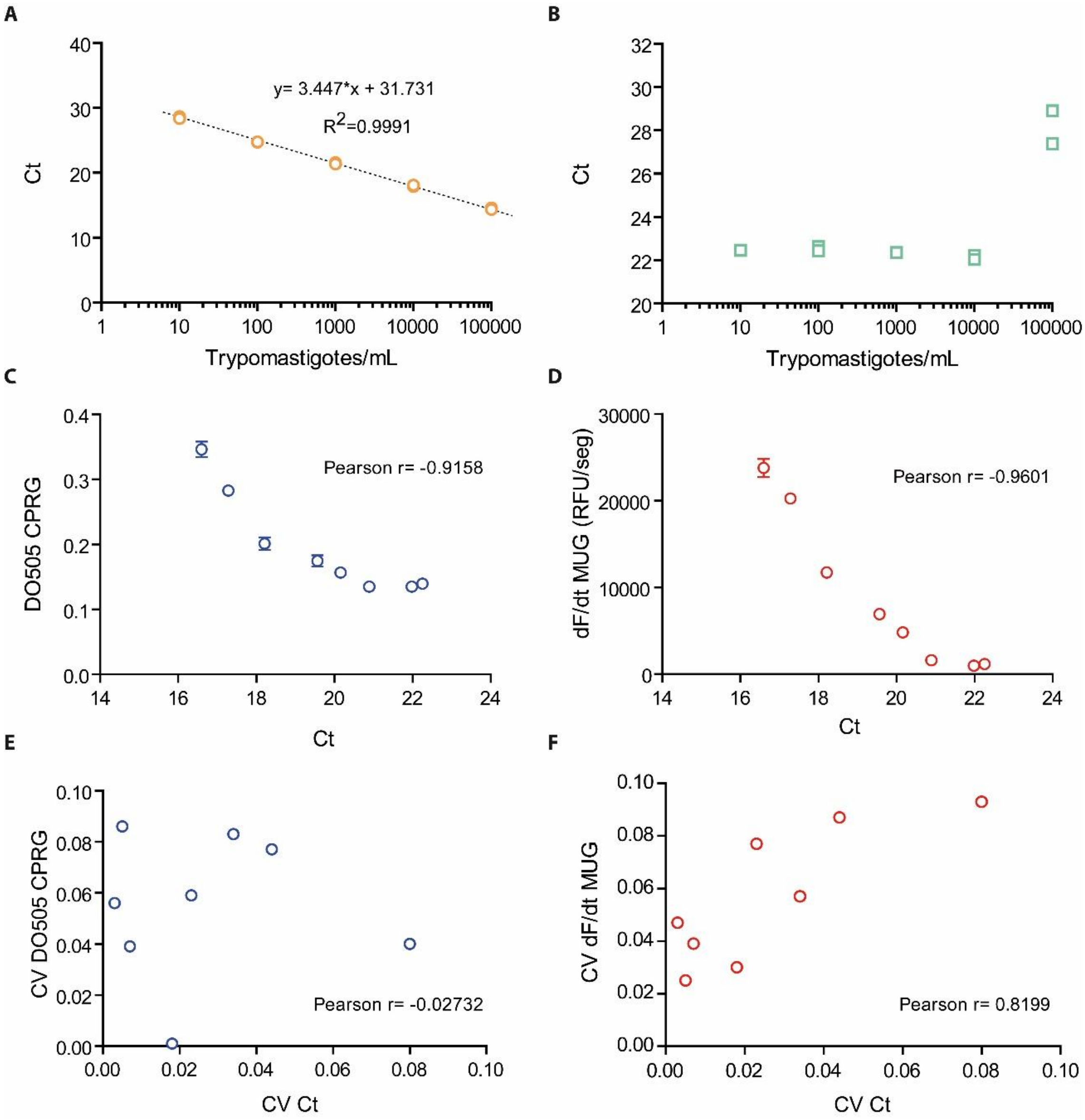


**Supplementary Figure S4. Analysis of parasite loads estimated by qPCR, their inter-replicate variability, and their correlation with those from enzymatic methodologies.** The amplification of SatDNA and Internal Amplification Control (IAC) targets was simultaneously monitored on different channels: A) Standard curve for the quantification of *T. cruzi* DNA by SatDNA amplification (range: $10\text{-}10^5$ trypomastigotes/mL). The best parameters from the fitting of experimental data to the linear model are indicated. B) Amplification of the IAC target in the standard curve. Except for the highest parasite load, where delayed amplification was observed due to competition with the main reaction, Ct values remained around 22. In both panels, replicates are shown individually. Correlation analysis between β-galactosidase activity and the amount of SatDNA: C) Plot of $DO_{505}$ vs. Ct for the end-point assay. D) Plot of dF/dt vs. Ct for the continuous assay. Correlation analysis between the coefficient of variations (CV) from enzymatic methodologies and qPCR: E) CV of end-point assay (CV $DO_{505}$) vs. CV of qPCR (CV Ct). F) CV of continuous assay (CV dF/dt) vs. CV of qPCR (CV Ct). In all cases, the estimated Pearson r values are indicated.

**Supplementary Table SII. Summary of parasite loads estimated by qPCR in three replicates of infected cultures treated with increasing benznidazole concentrations.**

| Sample | Replicate | Ct_SatDNA | Qty (eq.par./ml) |
|---|---|---|---|
| BNZ 20 µM | 1 | 22.64 | 432.76 |
| | 2 | 22.43 | 500.19 |
| | 3 | 20.88 | 1,409.76 |
| BNZ 10 µM | 1 | 22.32 | 538.64 |
| | 2 | 22.21 | 577.25 |
| | 3 | 22.23 | 571.59 |
| BNZ 5 µM | 1 | 21.21 | 1,125.12 |
| | 2 | 19.07 | 4,704.60 |
| | 3 | 22.38 | 515.59 |
| BNZ 2.5 µM | 1 | 19.99 | 2,550.49 |
| | 2 | 20.23 | 2,164.98 |
| | 3 | 20.25 | 2,144.15 |
| BNZ 1.25 µM | 1 | 19.61 | 3,289.71 |
| | 2 | 19.45 | 3,665.85 |
| | 3 | 19.64 | 3,220.60 |
| BNZ 0.63 µM | 1 | 18.25 | 8,170.78 |
| | 2 | 17.57 | 12,799.53 |
| | 3 | 18.82 | 5,566.55 |
| BNZ 0.31 µM | 1 | 17.14 | 17,074.16 |
| | 2 | 17.07 | 17,906.42 |
| | 3 | 17.64 | 12,230.66 |
| BNZ 0.16 µM | 1 | 17.02 | 18,525.89 |
| | 2 | 16.52 | 25,963.87 |
| | 3 | 16.26 | 30,778.25 |

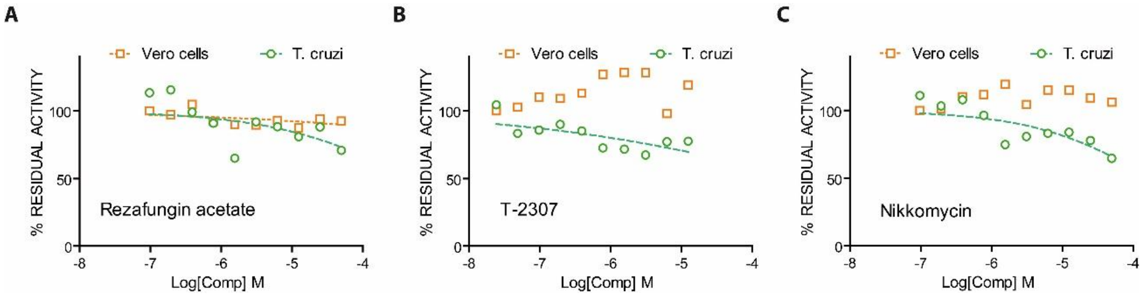


**Supplementary Figure S5. Representative dose-response curves for the antiparasitic and cytotoxic activities of inactive antifungals**. Antiparasitic activity against *T. cruzi* is represented with green circles, and the cytotoxic activity on Vero cells with orange squares. A) Rezafungin acetate. B) T-2307. C) Nikkomycin Z.

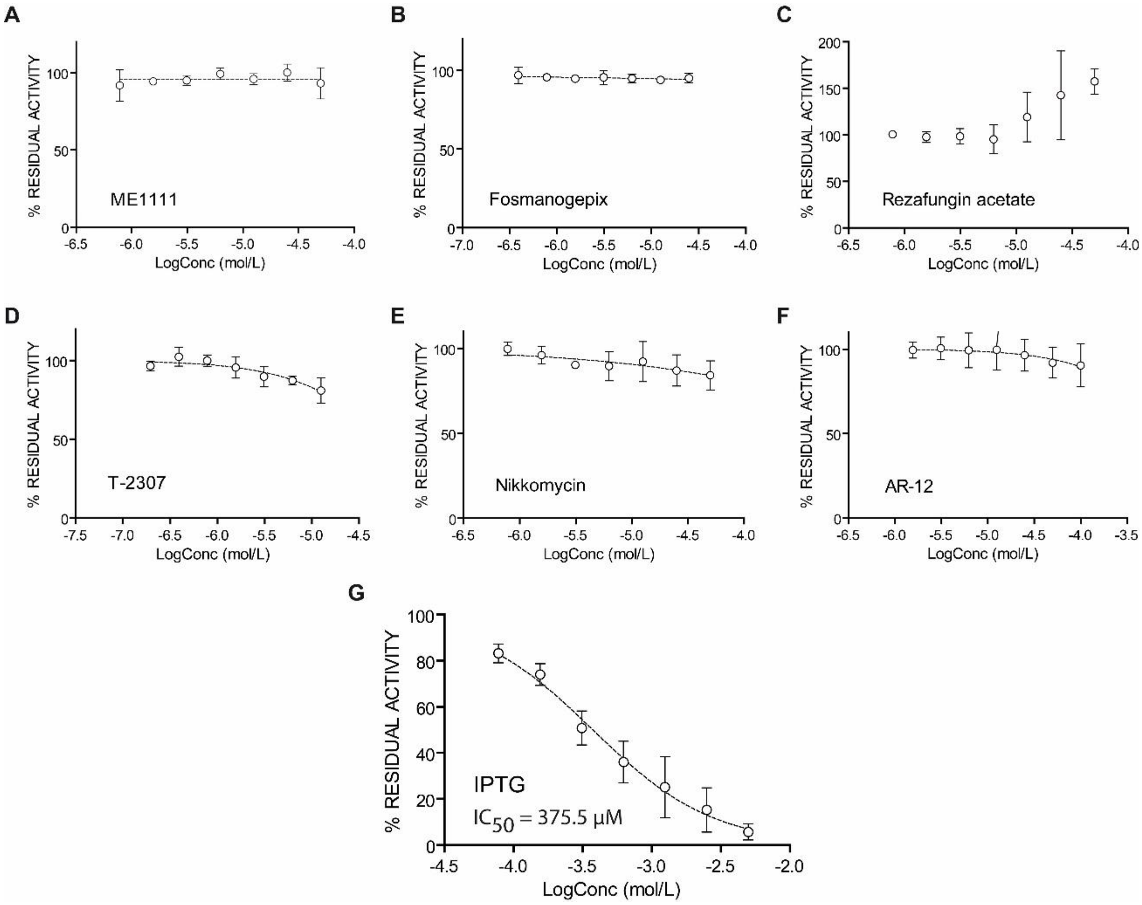


**Supplementary Figure S6. Analysis of β-galactosidase inhibition by antifungal compounds.** Increasing concentrations of investigational compounds were preincubated with a lysate of TulI β-gal trypomastigotes (dilution 1:2,000) in activity buffer (PBS supplemented with 2% RPMI, 0.25% DMSO, 0.5% NP-40, pH 7.4) before the addition of MUG substrate (10 µM). Steady-state slopes were derived from progression curves and used to build dose-response curves for each compound: A) ME1111. B) Fosmanogepix. C) Rezafungin acetate. D) T-2307. E) Nykkomycin Z. F) AR-12. G) Isopropyl β-D-1-thiogalactopyranoside (IPTG, positive control). Each data point represents the average of three replicates.

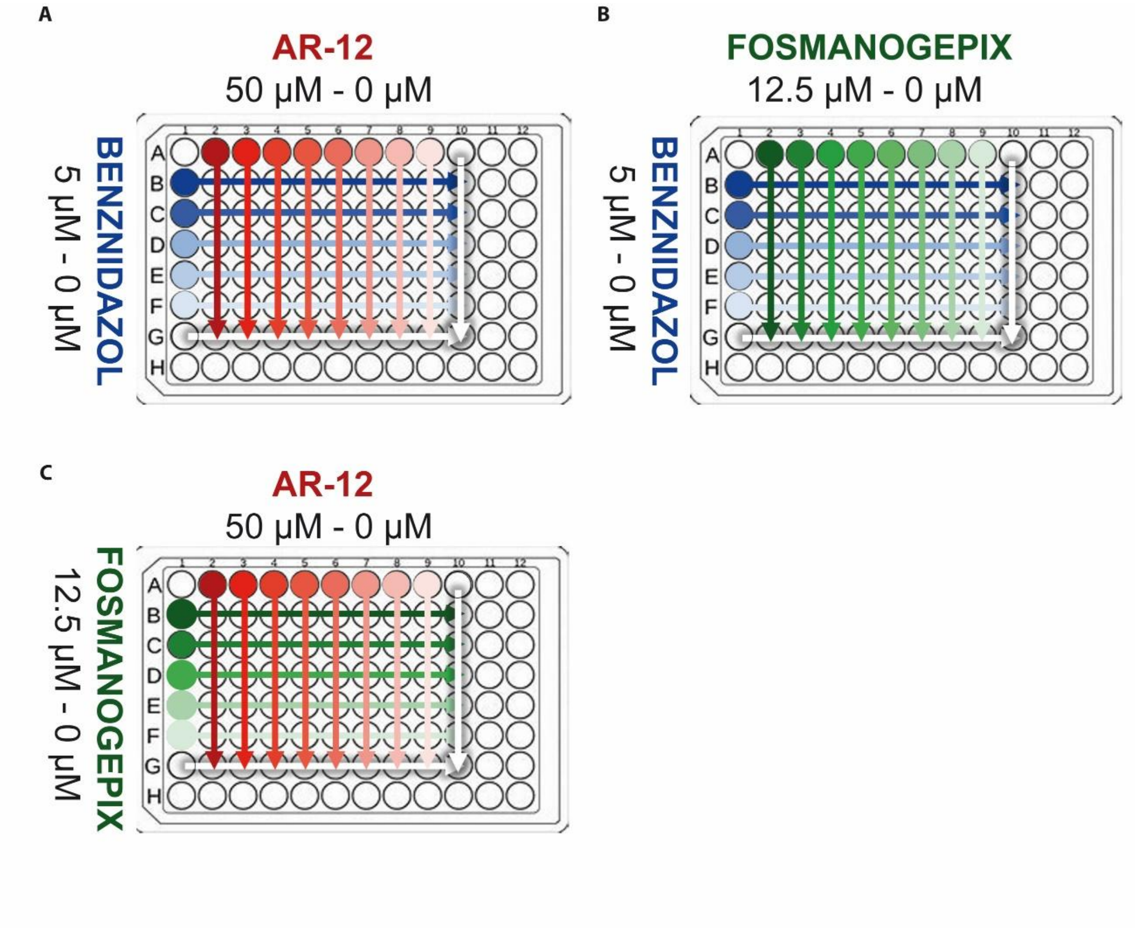


**Supplementary Figure S7. Plate diagram for the analysis of drug combinations with the semi-automatic continuous β-galactosidase methodology**. Three-fold serial dilutions of each compound in DMSO were performed by the OT-2 robot on the first column and row, as indicated. From these primary series, a combination matrix was formed by mixing 5 µL of each drug solution at the corresponding concentration. One microliter of each drug combination was added to 200 µL of media. After mixing, 100 µL (final DMSO concentration 0.5%) were incubated for 72 hours with infected Vero cells, as indicated in Materials and Methods. The final concentration of each drug in the culture ranged from 50-0 µM (AR-12), 12.5-0 µM (Fosmanogepix), and 5-0 µM (BNZ). In the diagrams, each drug is represented in a different color (AR-12: red, Fosmanogepix: green, BNZ: blue), and its concentration is indicated with color intensity. DMSO is indicated with white arrows (row G and column #10).
